# The prostate cancer therapy enzalutamide compared with abiraterone acetate/prednisone impacts motivation for exploration, spatial learning and alters dopaminergic transmission in aged castrated mice

**DOI:** 10.1101/2021.04.01.438009

**Authors:** Celeste Nicola, Martine Dubois, Cynthia Campart, Tareq Al Sagheer, Laurence Desrues, Damien Schapman, Ludovic Galas, Marie Lange, Florence Joly, Hélène Castel

**Affiliations:** Normandie Univ, UNIROUEN, INSERM, DC2N, 76000 Rouen, France, Institute for Research and Innovation in Biomedicine (IRIB), 76000 Rouen, France; Cancer and cognition Platform, Ligue Nationale contre le Cancer, 14000 Caen, France; Normandie University, UNIROUEN, INSERM, PRIMACEN, Rouen, France.; Clinical Research Department, Centre François Baclesse, 14000 Caen, France; Normandie Univ, UNICAEN, INSERM, ANTICIPE, 14000 Caen, France; University Hospital of Caen, 14000 Caen, France

**Author notes:** Co-first authors, these two authors contributed equally to this work. Corresponding author: Hélène Castel, INSERM U1239, Laboratory of Neuronal and Neuroendocrine Communication and Differentiation, University of Rouen Normandy, 25 Rue Lucien Tesnière, 76821 Mont-Saint-Aignan.

**Keywords:** metastatic resistant prostate cancer, enzalutamide, abiraterone acetate/prednisone, aged castrated mice, behavior, dopaminergic pathways

## Abstract

Cognitive side effects after cancer treatment, threatening quality of life (QoL) and adherence to treatments, now constitute a major challenge in oncology. Abiraterone acetate plus prednisone (AAP) and enzalutamide (ENZ) are next generation therapy (NGT) administered with androgen deprivation therapy to metastatic castration-resistant prostate cancer (mCRPC) patients. NGT significantly improved mCRPC overall survival but neurological side effects such as fatigue and cognitive impairment have been recently reported. We developed a behavioral 17 months-aged and castrated mouse model receiving *per os* AAP or ENZ during 5 days per week for six consecutive weeks. After behavioral tests, brain and plasma were collected for immunohistochemical studies. The objective was to elucidate the impact of NGT on spontaneous activity, cognitive functions and emotional reactivity, as well as neurobiological functions. ENZ exposure reduced spontaneous activity and exploratory behavior associated with a decreased tyrosine hydroxylase (TH)-dopaminergic activity in the substantia nigra pars compacta and the ventral tegmental area. A decrease in TH^+^-DA afferent fibers and Phospho-DARPP32-related dopaminergic neuronal activities in the striatum and the ventral hippocampus, highlighted ENZ-induced dopaminergic regulation whithin the nigrostriatal and mesolimbocortical pathways. ENZ and AAP treatments did not substantially modify spatial learning and memory or behavioral flexibility performances, but ENZ led to a thygmotaxis behavior impacting the cognitive score, and reduced c-fos-related activity of NeuN^+^-neurons in the dorsal hippocampus. These results establish the consequences of the mCRPC treatment ENZ in aged castrated mouse motivation to exploration and cognition, of particular importance for future management of patients elderly postrate cancer patients and their QoL.

## INTRODUCTION

It is now accepted that cancer treatments including brain irradiation and/or systemic chemotherapy induce cognitive dysfunctions including learning, memory, information processing speed and executive functions associated with brain structural and functional altered activity (1–3), all referred as “cancer related-cognitive impairment” or CRCI (4, 5). Long-term CRCI now constitutes a challenge for patients/survivors of localized non-central nervous system (CNS) tumors and may also be associated with targeted therapy such as those involving anti-angiogenic agents (6), novel immunotherapy (7) as well as new generation hormone therapy NGT (6–9). Currently the understanding of vulnerability factors of CRCI, most deleterious new cancer drugs and of biological-physiological mechanisms affecting patient neurological functions, fatigue or mood disorders represents important issues in terms of survival and quality of life (QoL) in particular in elderly cancer patients.

Prostate cancer (PC) is a major public health concern, being the second most common cancer among the leading causes of cancer-related death in men in the US/European occidental countries (10, 11), and represents the most frequent cancer in elderly population. In case of radical prostatectomy and/or radiation therapy following biochemical recurrence, a pharmacological androgen deprivation therapy (ADT) to abolish the gonadal testosterone (T) synthesis is proposed (12). When progression to metastatic castration-resistant prostate cancer (mCRPC) appears inevitable (13), NGT are now proposed to this older patient population (14). One consists in targeting the extragonadal and intratumoral cytochrome P450-17 (CYP17A1) enzyme activity converting pregnenolone and progesterone into T precursors, by abiraterone acetate, a prodrug of abiraterone, in association with prednisone (AAP), to avoid cortisol deficiency (14). The other strategy consists in the next-generation of androgen receptor (AR) irreversible high affinity antagonist enzalutamide (ENZ), impairing AR translocation to the nucleus and interaction with DNA androgen-response elements (15, 16).

These treatments significantly improve the overall survival (OS) of patients with mCRPC before or after chemotherapy (14, 17–19). For now, in the absence of better mCRPC therapy, the relative efficacy associated with good tolerability profile of AAP and ENZ (20) tends to increase their availability to elderly patients (21). Because aging by itself is associated with functional decline, this population of cancer patients appears at greater risk for increased age-related brain changes secondary to cancer and cancer treatments (22) impacting QoL. Interestingly, delayed health-related QoL alteration and pain progression was observed for ENZ or AA compared with chemotherapy (15, 23, 24), but their impact has never been evaluated in mCRPC patients on objective cognitive functions, despite a current clinical trial (21). Recently, neurological events were described with ENZ (25), confirmed in a prospective phase IV study showing more fatigue and subjective cognitive impairments in ENZ-than in AAP-treated mCRPC patients (26). These symptoms may be explained by central production of T and dihydrotestosterone (DHT) *via* peripheral steroid intermediates metabolism or *de novo* central synthesis by steroidogenic enzymes (27, 28), expressed in different brain areas (29) including hippocampal CA1-CA3 and dentate gyrus (DG) in rodent brain (30, 31). Together, NGT may impact brain functioning and induce fatigue related to QoL, as well as cognitive impairment in mCRPC elderly vulnerable patients.

The study herein aims at dissecting the direct brain impact of NGT in the context of aging and androgen deprivation in a behavioral preclinical model of aged castrated male mice. We explored the impact of oral treatment of ENZ or AAP on spontaneous activity, emotional reactivity including anxiety- and depressive-like behaviors, as well as spatial and learning memory. We found that ENZ exposure reduces spontaneous activity and exploratory behavior and impairs dopamine (DA) neurons of the substantia nigra pars compacta (SNpc) and the ventral tegmental area (VTA), leading to reduced DA production throughout nigrostriatal and mesolimbocortical pathways. Also, neither ENZ nor AAP treatments drastically altered spatial learning, memory and behavioral flexibility, but ENZ led to deficits in spatial cognitive strategies and to a decreased dorsal hippocampal (dHP) neuronal activity.

## METHODS AND MATERIALS

### Animal housing and management

C57Bl/6J Rj male mice of 15 months old were surgically castrated under the following institutional animal care and european guideline within the Janvier laboratory (Le Genest-Saint-Isle, France) following a conventional protocol. The mice were anesthetized, the surgical area was asyptically treated, the peritoneum incised and the left or right testicular fat pad holding the testis were gripped and cut using a steril scalpel. After cauterization of the testicular artery, the peritoneum is sutured, the skin is stapled with surgical wound clips and analgesia is performed to manage pain (carried by Janvier Labsoratories (Le Genest-Saint-Isles, France). At the age of 17 months, castrated mice were delivered in our institute animal facility. Mice were group-housed under controlled standard environmental conditions: 22 ± 1°C, under an inverse 12/12h light/dark cycle (light on: 00:00 to 12:00), with water and food available *ad libitum*. After 2 weeks of adaptation, animals were daily handled for weight monitoring during 1 week, and then throughout the treatment period. Treatment administration began at 17 months and 2 weeks of age while entire experimental sessions run for 6 additional weeks. The behavioral tests were performed during the active phase of mice (from 01:00 PM). For some control experiments, control aged non castrated mice were also obtained from Janvier Labs and tested at 18 months and 2 weeks aged on some behavioral tasks. All mice were euthanized at 19 months. The number and the suffering of animals were minimized in accordance with the guidelines of the European Parliament and Council Directive (2010/63/EU) and ARRIVE (Animal Research: Reporting of In Vivo Experiments) 2.0 guidelines (89). This project was approved by the “*Comité d’Ethique Normandie en Matière d’Expérimentation Animale*” and the French Research Minister (#7866-2016112115226170 v5) and carried out under the supervision of authorized operators (HC and MD).

### Treatment administration

For chronic treatments, mice received per os vehicle or NGT every day during 5 days per week for six consecutive weeks. 100 μL/10g/day of vehicle of ENZ (Veh-ENZ): 1% CMC (carboxymethylcellulose sodium), 1% tween 80, 5% dimethyl sulfoxide (DMSO) (Sigma Aldrich, Saint-Quentin-Fallavier, France) or ENZ (MedChem Express, Monmouth Junction, NJ, USA): 30 mg/kg in Veh-ENZ, vehicle of abiraterone acetetate (Veh-AAP): 20% hydroxypropyl β cyclodextrine (Hpβ CD), 0.037N HCl, 0.13X PBS in dH_2_O (Sigma-Aldrich) or abiraterone acetate-prednisone (AAP): 200 mg/kg of abiraterone acetate (Janssen Research & Development, Issy-les-Moulineaux, France) + 500 µg/kg prednisone (Sigma-Aldrich) Veh-AAP, every day during 5 days per week for six consecutive weeks. The treatments were administred *per os* with 22G feeding needles (Fine Science Tools, Heidelberg, Germany). Concerning the experimental session of Veh-AAP and AAP administration, food deprivation was done for all mice 2h before and 1 h after treatment because the composition of meals can affect AAP pharmacokinetic. Aged control mice were non castrated, did not receive oral administration of vehicle or treatment.

### Behavioral tests

Behavioral test panels were selected to evaluate the impact of the castration process (aged control mice, qualitative observational study) and of the treatment with ENZ or AAP versus vehicles on activity and exploration, emotional reactivity and cognitive functions. Thus, as soon as received, animals were distributed at random in the different experimental groups, at a rate of 5 mice *per* cage. All tests were performed as previously described (5) with slight modifications. The spontaneous activity and exploration of animals was evaluated in the open-field test (OFT). Anxious like behaviors were assessed by means of the elevated plus maze (EPM) and the light and dark emergence test (LDB). Depressive-like behaviors were evaluated with the tail suspension test (TST) and the forced swim test (FST). Then, the spatial learning, spatial memory as well as behavioral flexibility were measured in the Morris water maze (MWM). In total 5 behavioral experimental sessions were conducted. At the end of experimental sessions, brains and blood samples were collected from each group to lead immunohistochemical experiments on CNS slices and quantification of cytokine and hormones from plasma collection, respectively.

#### Open Field test

The general activity was measured during 10 min by placing mice in an OFT apparatus (45×45×31 cm, LxlxH). The OFT is a widely used device measuring exploratory and locomotion activity in a novel environment, a dopamine-dependent process (35). The animal was placed into the center of the apparatus and several behavioral parameters (distance crossed, vertical activity and grooming) were automatically analyzed by a the videotracking system Anymaze (Stoelting®, Dublin, Ireland). Rodents typically prefer not to be in the center, lit area of the apparatus and tend to walk close to the walls (thigmotaxis) since this is a novel, and presumably, stressful, environment to the animal. So distance and time spent into the center part of the device is an indication of anxiety (90), but motor activity and exploration can also provide a behavioral incentive value (91).

#### Elevated plus-maze

Anxious like behaviours were evaluated by means of the EPM during a unique session of 5 min as previously described to evaluate state anxiety (*i.e.* in a forced situation) (92) and by LDB (supplementary information). It allows to determine emotional reactivity of animals through a conflict between secure parts of the maze (2 enclosed arms) and aversive parts of the maze (2 open arms) of a device elevated at 41.5 cm above the floor. Time spent and number of entries into the open and enclosed arms of the maze were measured and the percent number of entries and percent time in open arms were calculated.

#### Tail suspension test and Forced swim test

Depressive-like behaviors were evaluated in the TST (93) or the FST (94), two standard procedures used to evaluate the antidepressant activity of pharmacological compounds. In TST, mice were suspended at a height of 20 cm above the floor and are surrounded by a enclosure and the total duration of immobility (passive hanging) between periods of wriggling to avoid aversive situation was measured during a period of 6 min, and the latency to the first immobility was also noted, as previously described (95). In FST, mice were placed during 6 min into a cylinder (17 cm in diameter) filled with 25°C tap water (at a height of 13 cm) with no way out. The immobility duration (excluding movement necessary to keep the head above water or to float) was measured as an indication of the behavioral despair of mice (95).

#### Morris water maze

Spatial learning and memory performances, as well as learning plasticity, were evaluated in the MWM (95, 96) as previously described. Briefly, mice placed in a water tank and have to escape water by finding a hidden platform (9.7 cm in diameter). Behavior of animals was videotracked with Anymaze (Stoelting®, Dublin, Ireland). Tests were conducted during several sessions (95), each session being composed of of 4 trials of 60 s maximum each (except the probe test). The first session (familiarization) was conducted by placing the escape plateform in the center of the pool. Mice were placed on it during 20 s and immediately after, the mice was placed in water, facing the wall, at one of the 4 starting points (Est, E; North, N, West, W; south, S) for a trial of maximum 60 s. If mice did not find the escape plateform within the 60 s maximum duration of trial, animals were manually placed on it during 20 sec. Then 3 successive trials, with an intertrial intervall of 30 min, were conducted with a departure from one of the 3 other starting points. During the 4 consecutive days, the spatial learning abilities were tested by placing the escape platform in the NW quadrant. Animals had to use the distal visual cues around the basin to find the platform location. Each day, 4 trials of 60 s maximum were conducted. The 4^th^ day of the hidden platform phase, 2 h after the end of the 4^th^ trial, a probe test was conducted by removing the platform and allowing animals to swim into the pool for a unique trial of 60 s. Time spent by mice in the previously correct quadrant (NW) was measured. Spatial memory abilities were evaluated at the 3^th^ day, during the retrieval test. The platform was hidden in the NW quadrant and mice had a maximum of 60 s to find its location, for four successive trials for a unique day. In the 9^th^ day, the platform was moved from the NW to the SE quadrant and behavioral flexibility was studied and the procedure was the same as previously described for the hidden platform test.

### Swim search strategy analysis, dataset preparation and cognitive score from MWM

In MWM spatial learning session, trial after trial the animals accumulate knowledge of the spatial relations of the cues and the specific contexts allowing effective navigating routes to find the hidden platform. Sterotypic sequences of search patterns with swim path trajectories can be analyzed as they can be considered representative of cognitive functions during the task. Graziano et al. (72), put forward automatization of several explorative strategies. Here, as previously established (97), image files corresponding to tracking trajectories for each mouse in all trials along the 4 consecutive training days were obtained (AnyMaze; Stoelting Co., Wood Dale, IL) and manually labelled according to the 6 categories of trajectories as previously described (97): thigmotaxis, scanning, circling, focal search, rotating and direct swim. If some traits were mixed within one trial, most prominent trajectory was adopted as label. Representative trajectories are shown in Fig 4D as previously described (97), a convolutional neural network (CNN) with two convolution layers were used to classify the swim path trajectories, by means of the minimal available program from the repository (https://figshare.com/s/90d7b2d038551efe08ec) (97) which help us designing our study model. In this work, recognition model output was then use to assign each image to the respective subset. For the model construction, swim path images of familiarization, retrieval and flexibility phases were collected for each mouse group from AnyMaze (Stoelting®, Dublin, Ireland). The dataset (420 images) was divided into two sub-datasets; 80% of the data were used for the training stage and the other 20% were used for validation. In this model proposed by Higaki et al. (97) we integrated new instructions to create an unbiased automatic tool able to recognize unlabelled images. Thus, unlabelled swim path images of learning phase were collected and randomly passed in our modified model. The original format was a Microsoft Windows Bitmap Image (BMP) file with 140×120 pixel size and 32-bit color data. All image data were converted to grayscale pictures of reduced pixel size (48 x 48) using a free image processing tool of Python interpreter (Pillow; Alex Clark and Contributors). As previously (97), pixel values derived from each image file were divided by 255 for standardization and passed to our modified CNM as input data. Whole image files were randomly rearranged before being assigned to each subdataset. Repeated holdout cross-validation was performed 5 times for each randomly rearranged dataset, and the average score was employed as the valid outcome. All these modeling processes were provided by Chainer, previously validated for this procedure (97, 98). In addition, we built a score corresponding to cognitive performance in each training trial as previously described (46, 97), attributing a higher score to the best efficient cognitive strategy: Thigmotaxis: 0, Scanning: 1, Circling: 2, Focal search: 3, Rotating: 4, Direct swim: 5. Average cognitive score within 4 trials and during 4 learning days in each mouse was calculated and used for the analysis.

### Brain analyses

At the end of the treatment sessions mice were subjected to intraperitoneal injections of BrdU (5-bromo-2-deoxyuridine, Sigma-Aldrich) at 50 mg/kg once a day during 4 days to label adult-generated progenitor cells in the DG of the hippocampus, and dopaminergic activity in the SNpc and VTA and consequences via afferences in the striatum and ventral hippocampus was investigated by immunohistochemistry on brain slices. After the last injection of BrdU on the the fourth day, mice were anesthetized using isoflurane (Isovet, Osalia, Paris, France), decapitated and each brain was immediately removed and frozen in 2-methylbutane (Sigma-Aldrich) at −30°C and then stored at −80°C until use. Brain sections (20 μm thick) were cut with cryostat, from the anterior part of the dorsal hippocampus (anteroposterior, 1.08 mm from the Bregma −1.46 mm). Every 12^th^ section, each separated by 240 mm, was mounted on slides coated with gelatin-chrome alum (VWR International, Leuven, Belgium) and stored at − 20°C until processing. Eight hippocampal sections from 3 to 4 animals *per* group were stained simultaneously for immunohistochemical observations and quantifications. Slices were post-fixed with formaldehyde (4%, Sigma-Aldrich) for 15 min at 4 °C, rinsed in PBS (0.1 M, pH 7.4, Sigma-Aldrich), permeabilized with X-100 triton (Fisher Scientific, Illkirch, France) 0.05% and non-specific binding sites were blocked with a permeabilization/blocking solution containing PBS (Sigma-Aldrich), 0.5% BSA (Roche, MannHeim, Germany), 0.1% X-100 triton and 10% NDS (Normal Donkey Serum, Sigma-Aldrich) for 60 min at room temperature (RT). For BrdU immunostaining, brain slices were first incubated in a 2N-HCl for 45 minutes at 45°C and then rinsed in a boric acid (Sigma-Aldrich) solution 0.1 M at pH= 8.5 for 10 min. Brain slices were incubated overnight (12h) at 4°C with the primary antibodies of interest diluted in the permeabilizing/blocking buffer (PBS, 0.5% BSA, 0.1% X-100 triton and 1% NDS). The following primary antibodies and the corresponding dilutions were used as followed: anti-GFAP (Goat, Abcam, Cambridge, UK, AB53554,1:1000) or anti-GFAP (Rabbit, Dako, Agilent, Santa Clara, CA, USA, Z0334), anti-AR (Rabbit, Abcam AB74272, 1:300), anti-BDNF (Goat, St John’s Lab, London, UK, STJ73299, 1:1000), anti-TH (Goat, Abcam AB101853) or anti-TH (chicken, Abcam AB76442,1:500), anti-Phospho-DARPP-32 (P-DARPP-32, Rabbit, Bioss, Boston, MA, USA, bs-3118R, 1:200), anti-BrdU (Sheep, Abcam AB1893, 1:200), anti-NeuN (Rabbit, Abcam AB104225) or anti-NeuN (Chicken, MyBiosource MBS837654, 1:1000), anti-c-fos (Rabbit, Abcam AB190289), anti-DCX (Guinea pig, Millipore, Burlington, MA, USA), anti-D1R (Rabbit, Abcam, AB20066, 1:200) or anti-CYP17A1 (Rabbit, LifeSpan Biosciences Inc.,Seattle, WA, USA, LS-C352100, 1:100). The following day, sections were rinsed (PBS, 4×5 min) and incubated (2h, RT) with the appropriate secondary Alexa Fluor-conjugated antibodies diluted in the permeabilizing/blocking buffer (2h, RT). Secondary antibodies were used as followed: Alexa 647-donkey anti-goat (Abcam AB150131, 1:1000), Alexa 488-donkey anti-rabbit (Invitrogen, Carlsbad, CA, USA, A21206, 1:1000), Alexa 488-donkey anti-sheep (Invitrogen A11015, 1:1000), Alexa 488-donkey anti-rat (Invitrogen A31573, 1:1000), Alexa goat anti-guinea pig (Invitrogen A11073, 1:1000) and Alexa 488-goat anti-chicken (Abcam, AB150173). Sections were then rinsed with PBS (3×5 min at RT) and then cell nuclei were stained with a DAPI solution (Sigma-Aldrich, 1μg/mL) incubated for 5 min at RT. Slices were embedded in a Mowiol solution, dried overnight at RT, and then stored at 4°C until they were observed with an epifluorescence microscope (PRIMACEN, uprightt microscope Nikon 1, Nikon, Champigny/marne, France) as first evaluation, and by confocal microscopy (PRIMACEN, Upright confocal microscope, Leica Microsystems, Nanterre, France) for image acquisition and quantification. Labeled structures were assessed in 4 sections of 20 µm thickness (anteroposterior from bregma 0.62 mm, interaural 4.42 mm for striatum, at bregma − 1.82 mm, interaural 1.98 mm for dorsal hippocampus (dHP), at bregma −3.16 mm, interaural 0.64 mm for ventral hippocampus (vHP) at bregma −3.08 mm, interaural 0.72 mm for the substantia nigra pars compacta (SNpc) and ventral tegmental area (VTA) with SP8 confocal microscope (Leica, Heidelberg, Germany). The density, fluorescence intensity and total number of cells were quantified by using Image J software (version 1.8.0, NIH, USA).

### Statistical analysis

Prior to analysis, each treatment group normality was checked by Shapiro-Wilk normality tests, and variances were analyzed. The potential existence of outliers was verified thanks to the Grubs test. Based on the groups normality and analysis of variances, data were analyzed using either *(i)* two-way ANOVA followed by Sidak’s multiple comparisons test or t-test for *parametric analysis*, or *(ii)* Kruskal-Wallis followed by Dunn’s multiple comparison test or Mann-Whitney test for *non-parametric analysis*. For the analysis of swim path strategies in the Morris water maze, Chi-square test (χ^2^ test) with Yates’ continuity correction was used (Confidence interval 95%). For cognitive score analysis, statistical analysis was performed by repeated measures (RM) one-way ANOVA, both using the Tukey’s test for multiple comparisons. For all analyses, p value ≤0,05 was considered as significant. Elisa quantifications were performed from 6 to 9 plasma samples of each group. For immunohistochemistry analysis, based on groups normality and equivariance, data were analyzed using either *(i)* Mann-Whitney test for non parametric analysis, or *(ii)* unpaired t-test for parametric analysis. For all analyses, a P value ≤ 05 was considered as significant. Data are expressed as scatterplots and mean ± Standard Error of the Mean (SEM) and analyzed using GraphPad Prism-7 software (La Jolla, California, USA).

## RESULTS

We explored for the first time the neurological impact of NGT in aged castrated mice to model a clinical situation encountered by mCRPC patients first treated *via* ADT and then receiving either ENZ or AAP (Fig 1A). Our work is mainly focused on the potential regulatory mechanims of ENZ or abiraterone (AA), blocking the enzymatic activity of CYP17A1 in adrenal glands (and prostate) but also within some brain areas (Fig 1A). We chronically administered 100 µL/10g of ENZ or AAP *per os* during 6 weeks to 17.5 months old C57BL/6 male mice, castrated at 15 months old to mimick ADT. This model is based on the assumption that CYP17A1 expressed in brain structures is likely responsible for the conversion of pregnenolone and progesterone towards T and potentially DHT in one hand, and in estrogens in the other hand (Fig 1B). This local androgen production may control AR-activating signaling pathways in selective brain areas, regulating aged mouse neurobehavioral responses in the context of androgen deprivation.

**Fig 1.**
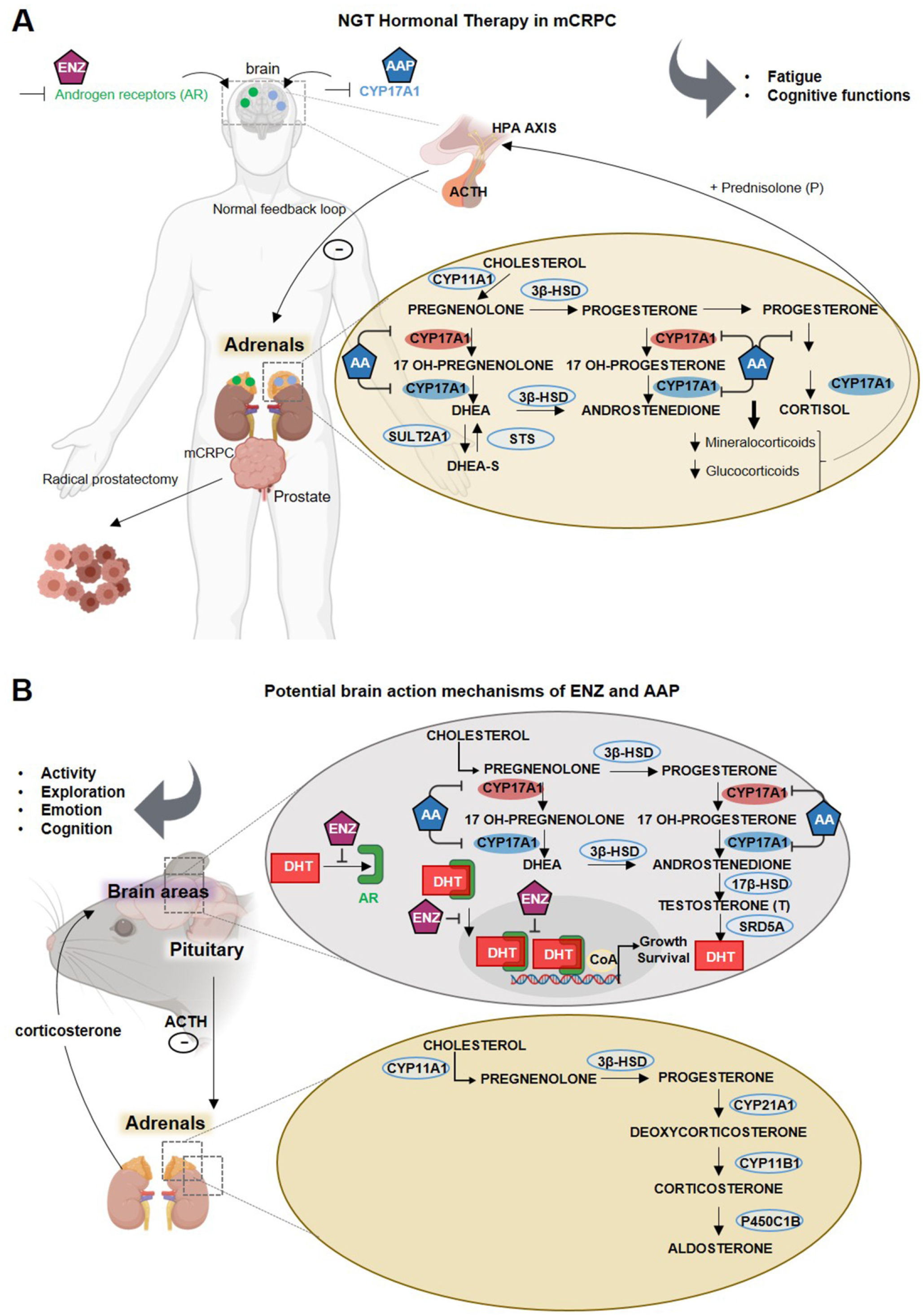
New generation therapy targeting androgen biosynthesis in addition to androgen deprivation therapy against metastatic castration-resistant prostate cancer: potential mechanisms of action on brain function using a mouse model. (A) Schematic of enzalutamide (ENZ) and abiraterone acetate-prednisone (AAP) as treatment options for metastatic castration-resistant prostate cancer patients and of their sites of action such as prostate tissue, adrenals and potentially the brain. Cytochrome p450c17A1 (CYP17A1), an ER-localized enzyme expressed in both prostate and adrenal glands, and potentially in human brain, plays a critical role in the androgen biosynthesis pathway. In the adrenals, CYP17A1 catalyzes two critical steps through its 17α-hydroxylase and C17,20-lyase activities. The 17α-hydroxylase activity of CYP17A1 converts pregnenolone to 17α-pregnenolone and progesterone to 17α progesterone. The C17,20-lyase activity of CYP17A1 converts 17α-hydroxypregnenolone to dehydroepiandrosterone (DHEA) and 17 α -hydroxyprogesterone to androstenedione, which are subsequently converted to testosterone and/or 5α dihydrotestosterone (DHT) by other steroidogenic enzymes. After radical prostatectomy and androgen deprivation therapy, androgens can be mainly synthesized from cholesterol through multiple enzymatic reactions within adrenals. Abiraterone is a novel CYP17A1 inhibitor that acts to inhibit its 17α-hydroxylase and/or C17,20-lyase activities. To ensure a normal mineralcorticoid feedback loop, low doses of prednisone are co-administered with AA in order to suppress ACTH, thereby restoring normal cortisol physiology in patients. ENZ is an irreversible antagonist with high affinity for the AR expressed both in adrenal glands and also within the brain. (B) NGT Hormonal therapy potential action mechanisms on mouse brain. Representation of ENZ and AAP potential action mechanism in castrated mice illustrating the potential binding of ENZ on AR in the rodent adrenals but also in the brain or the inhibitory action of AAP on CYP17A1 expressed in adrenals and in rodent brain. AA may impact stereroidogenesis within adrenals showing the cholesterol transformation into pregnenolone by Cytochrome P450 Family 11 Subfamily A Member 1 (CYP11A1), being metabolized by 3β-hydroxysteroid dehydrogenase (3 -HSD) into progesterone. Through multiple enzymatic reactions within adrenals progesterone will be transformed in the mineralcorticoid aldosterone. In the upper panel, representative scheme of the streroidogenesis within CNS showing the cholesterol transformation into pregnenolone by Cytochrome P450 Family 11 Subfamily A Member 1 (CYP11A1) in the inner mitochondrial membrane, being metabolized by 3β-hydroxysteroid dehydrogenase (3β-HSD) into progesterone. AAP, by blocking CYP17A1, may impair progesterone transformation into 17-OH-progesterone, 17-OH pregnenolone and Dehydroepiandrosterone (DHEA) as well as transformation of DHEA into androstenedione by 3β-HSD, transformation of androstenedione to testosterone by the 17β-hydroxysteroid dehydrogenase (17β-HSD) and reduction of T to DHT by 5α-reductase (SRD5A). ENZ treatment potentially impairs the binding of T/DHT to AR, the translocation of ligand-activated-AR into the nucleus and its action as transcription factor when associated with the coenzyme A (CoA). Modulation of AR signaling transduction activated by local synthesis of T or DHT, should thus regulate neurotransmission, neurogenesis, and brain plasticity and selective behavioral responses as activity, emotion, exploration or cognitive functions. 17β-HSD, 17β-hydroxysteroid dehydrogenase; 3β-HSD, 3β-hydroxysteroid dehydrogenase; AAP, abiraterone acetate-prednisone; ACTH, Adrenocorticotropic hormone; AR, androgen receptor; CNS, central nervous system; CoA, coenzyme A; CYP11A1, Cytochrome P450 Family 11 Subfamily A Member 1; CYP17A1, cytochrome P450 family 17 subfamily A member 17; DHEA, dehydroepiandrosterone; DHT, dihydrotestosterone; ENZ, enzalutamide; ER, estrogen receptor; NGT, new generation hormone therapy; SRD5A, 5α-reductase.

### ENZ but not AAP treatment reduced spontaneous activity, exploration and motivation to effort with no impact on plasma cytokines

We evaluated the behavioral consequences of *per os* ENZ or AAP treatment during 5 consecutive days for 6 weeks, on general spontaneous activity and exploration, emotional reactivity and spatial learning and memory in aged castrated mice by means of validated behavioral tests (Fig 2A). No significant effect of ENZ or AAP were detected on the course of weight gain (S1 Fig). Based on the findings that low levels of T in men can be associated with an inflammatory status in different situations such as in healthy elderly population (32, 33) or hypogonadism (34), common cytokines were measured in plasma of aged castrated mice and non-castrated/non treated mice. We did not show a significant difference in the pro-inflammatory, pluripotent, chemotactic or anti-inflammatory cytokine levels between ENZ or AAP and their vehicle groups, and the control group (S1 Table), discarding a potential consequence of NGT on systemic cytokine modifications in aged subjects.

**Fig 2.**
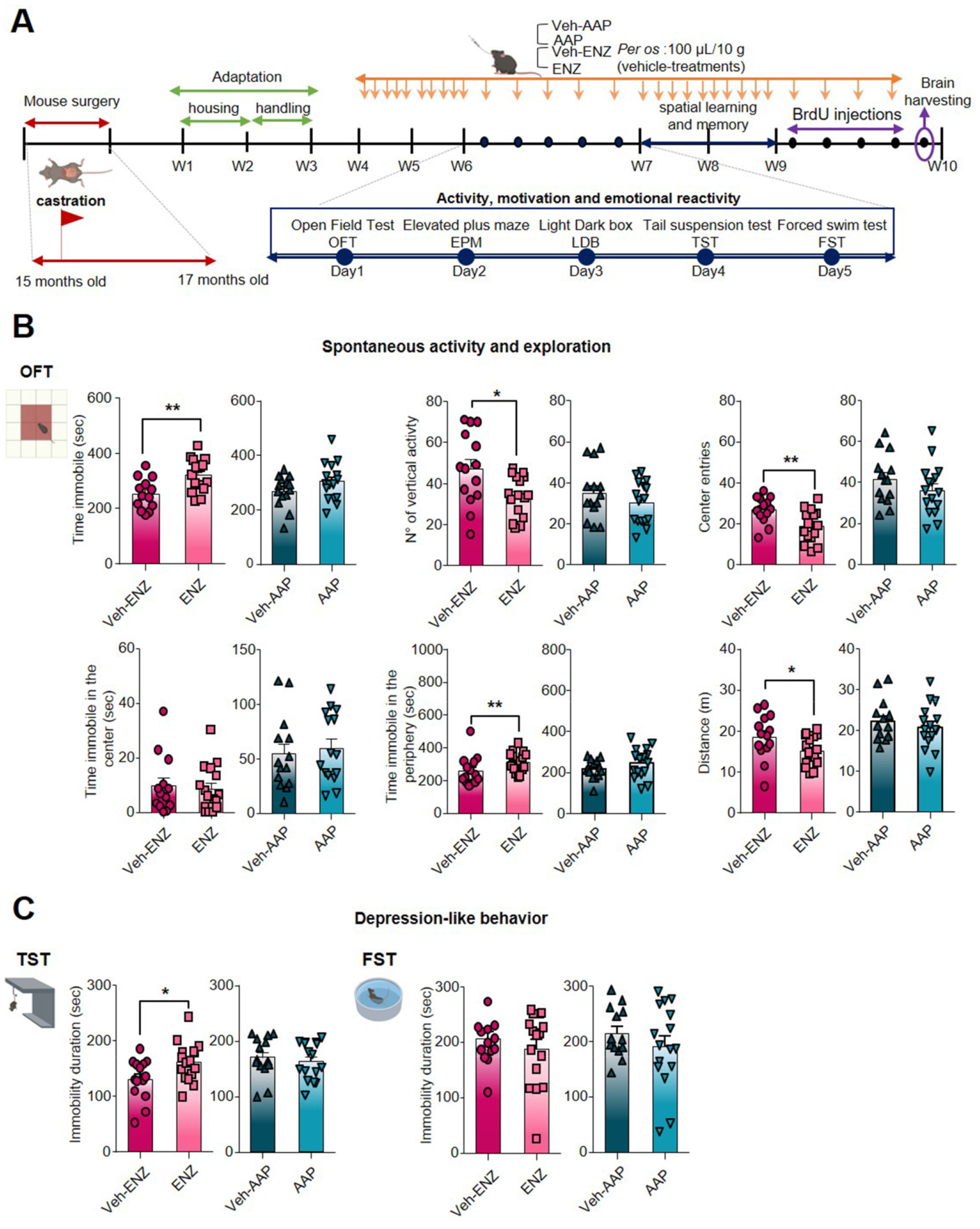
Characterization of ENZ or AAP impact on incentive exploratory behavior and motivation to effort of mice. (A) Schematic diagram showing the timeline for treatments and behavior experiments during ENZ and AAP treatments. C57BL/6J Rj mice were castrated at 15 months old, were housed under controlled environment from 17 months old, while treatments were administered orally (*per os*) once a day during 5 days a week, from week 4 (W4) to week 10 (W10), as indicated (orange arrow). Behavioral evaluations (blue arrow) started at W6, using open-field test (OFT) on D1, elevated plus maze (EPM) on D2, light dark box (LDB) on D3, tail suspension test (TST) on D4 and forced swim test (FST) D5. Mice brain and plasma were collected (purple arrows) at the end of W10 after the cognitive evaluation session. (B) and (C) Impact of ENZ or AAP compared with vehicle on spontaneous activity and exploration in OFT and on resigned behavior/motivation to effort in TST (left) and FST (right panel). Behavioral statistical analysis were performed between vehicle (*n* = 14) and treated (*n* = 16) mice using unpaired *t*-test or Mann-Whitney test; Bars are mean ± SEM while symbols individual data points; \**P*□≤□0.05, \*\**P*□≤□0.01. AAP, abiraterone acetate-prednisone; D, day; ENZ, enzalutamide; EPM, elevated plus maze; FST, forced swim test; LDB, light dark box; OFT, open-field test; TST, tail suspension test.

The impact of ENZ and AAP was tested by using behavioral paradigms for spontaneous activity and exploration in a new environment, anxiety, depressive-like behavior involving motivation to effort, all validated in non castrated aged mice (S2 Table). In OFT, ENZ-treated compared with Veh-ENZ groups showed a significant increase in time spent immobile, a decrease number of vertical episodes, center entries and distance crossed (Fig 2B and S3 Table), suggesting a reduction of locomotion for exploration. In contrast, AAP- and Veh-AAP-treated mice exhibited normal activity, locomotion and exploration, and no group effect was observed for main parameters, except a decreased grooming time (S3 Table), suggesting a reduced arousal in an aversive/stressing environment. Because mice exposed to EPM and LDB tests showed no alterations after ENZ or AAP compared with Vehicle treatments (S3 Table), even if AAP was associated with more head dips (S3 Table), it appears that anxiety-like behaviors are not worsen with NGT. To verify depressive-like behavior and/or lack of sustained expenditure to effort, resignation was assessed by presenting rodents FST and TST. In FST, neither ENZ- nor AAP-treated groups exhibited significant changes in immobility duration and latency to the first immobile episode (Fig 2C) compared with Veh-treated groups. However, in TST, ENZ mice exhibited an increase of time spent immobile compared with Veh-ENZ (Fig 2C), while no differences were detected in AAP compared with Veh-AAP mice (S3 Table).

### ENZ treatment altered the nigrostriatal network by inhibiting the tyrosine hydroxylase- and Phospho-DARPP32-related dopaminergic activities

Radar plots of Fig 3A highlight that ENZ diminished spontaneous locomotor activity and exploration whereas AAP remained without significant deleterious effects on emotional or spontaneous activity. Dopaminergic pathways have been shown to drive exploratory and locomotion behaviors promoting the positive valuation of novel stimuli as well as motivation to explore (35). In rodent brain, DA projections from SNpc to the striatum (Nigrostriatal pathway) are critical for activity or action sequence initiation and performance (36) while DA projections from VTA to the ventral hippocampus (vHP) (part of the mesolimbocortical pathway) are involved in novelty detection and motivation to explore (37, 38). Based on previous published evidence suggesting that AR can be expressed in SNpc and VTA (39, 40), we focused here on the consequences of ENZ or AAP treatment on DA-mediating signalings involved in locomotor activity and motivation to exploration (Fig 2A). We analyzed AR expression in a DA subpopulations co-stained with an anti-tyrosine hydroxylase (TH) antibody in both SNpc and VTA midbrain nuclei from brain slices of ENZ-, AAP- and Veh-treated groups. AR expression remained constant among the different groups, some neurons being co-stained with TH (Fig 3B). The number of TH^+^-cell/µm^2^ appeared significantly diminished in ENZ-treated groups compared with Veh-ENZ in VTA and SNpc, whereas no difference in TH expression level was found between AAP and Veh-AAP conditions (Fig 3D). Importantly, CYP17A1 appeared expressed in both SNpc and VTA areas of ENZ, AAP and respective Veh-treated groups (Fig 3C), indicating that components of local synthesis of T and DHT and of AR signalings are present in the midbrain DA neurons.

**Fig 3.**
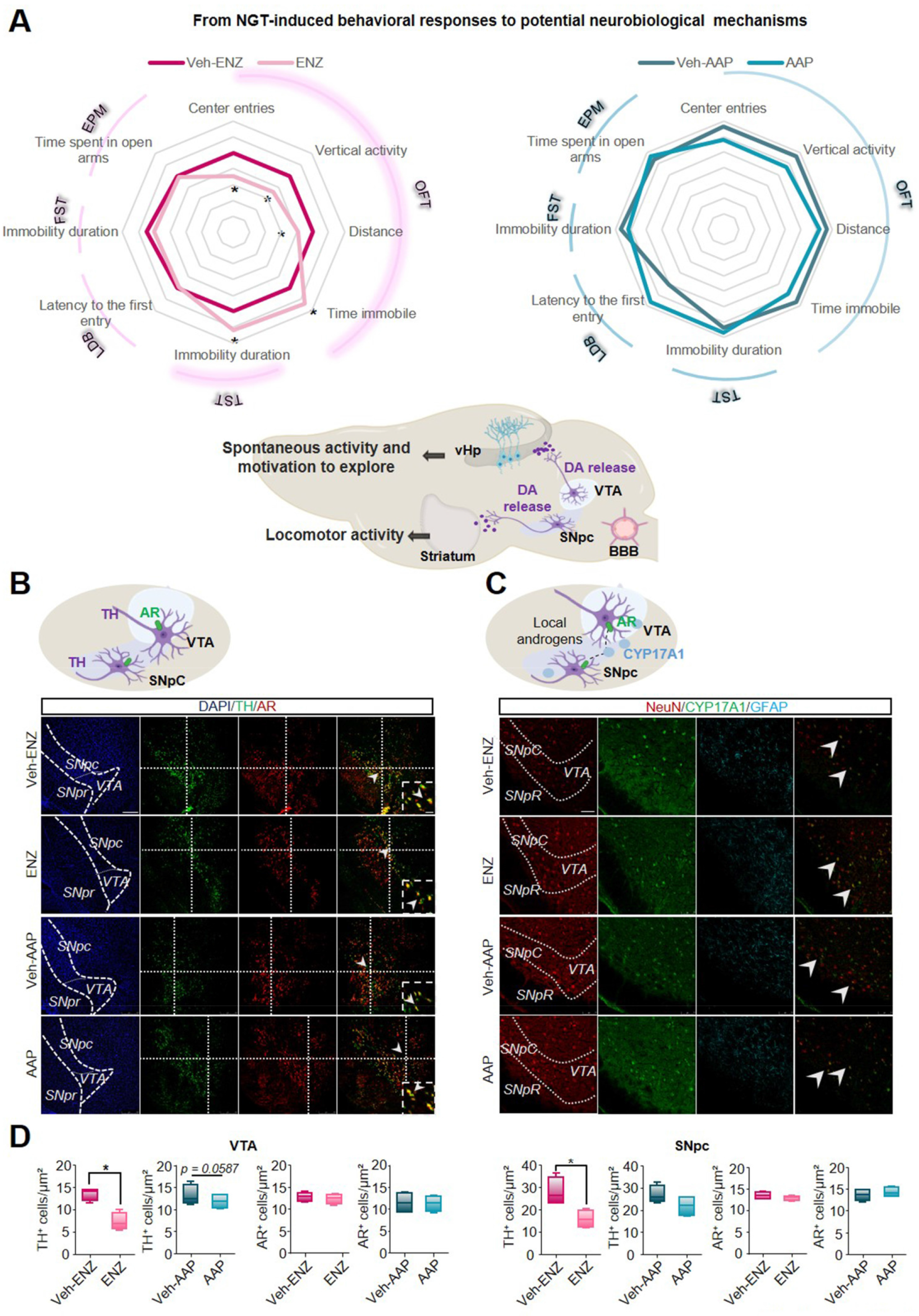
AR and CYP17A1 are expressed in the substantia nigra pars compacta and ventral tegmental area and ENZ inhibits dopaminergic activity in both midbrain nuclei. (A) Comparative Radar Chart of performance index scores across emotional and exploratory behavioral domains after ENZ (left panel) or AAP (right panel) treatments compared with Vehicles as presented in Fig 2A and Table S3. Based on the behavioral modifications, schematic representation (Middle panel) of nigrostriatal and corticolimbic circuits potentially involved in ENZ-altered locomotor activity and exploration. Based on previous studies, AR may be expressed in ventral tegmental area (VTA) and substantia nigra pars compacta (SNpc) dopaminergic neurons. SNpc projects dopamine (DA) axons to the striatum (nigrostriatal pathway) while VTA projects DA axons to ventral hippocampus (vHp) (corticolimbic pathway), also expressing AR. Statistical analysis was done on performances of vehicle (*n* = 14) and treated (*n* = 16) mice using unpaired *t*-test or Mann-Whitney test; Data are mean □±□SEM; \**P*□≤□0.05. (B) Schematic drawing of SNpc and VTA dopaminergic neurons expressing AR. Tyrosine hydroxylase (TH) (green) and AR (red)-immunoreactivity in the SNpc and VTA of ENZ-, AAP- and Vehicle-treated mice. The boxed areas showed magnification of clusters of TH^+^/AR^+^ cells. Scale bar: 250 μm and 100 μm. (C) Schematic drawing of Cytochrome P450 Family 17 Subfamily A Member 1 (CYP17A1) expression in SNpc and VTA and potential binding of local androgens on AR expressed by dopaminergic neurons. Representative example of CYP17A1 (green), NeuN (red) and GFAP (cyan) immunoreactivities in the SNpc and VTA of ENZ-, AAP- and Vehicle-treated mice. Scale bar: 250 μm. (D) Whisker boxes represent the number of TH cells and AR cells in VTA (left panel) and in SNpc (right panel) of ENZ- and AAP-treated mice compared with respective vehicles. Statistical quantification was performed by using Mann-Whitney test. Data are represented as box and whiskers and mean □±□SEM; (*n*□=□4 mice)\**p*□<□0.05. AAP, abiraterone acetate-prednisone; AR, androgen receptor; BBB, brain blood barrier; CYP17A1, cytochrome P450 family 17 subfamily A member 17; DA, dopamine; ENZ, enzalutamide; EPM, elevated plus maze; FST, forced swim test; GFAP, Glial Fibrillary Acidic Protein; LDB, light dark box; NeuN, Neuronal Nuclei Antigen; OFT, open-field test; SNpc, substantia nigra pars compacta; TH, Tyrosine hydroxylase; TST, tail suspension test; vHp, ventral hippocampus; VTA, ventral tegmental area.

### ENZ inhibited the DA-related P-DARPP-32 activity in D1R-expressing neurons of the striatum and the ventral hippocampus

To verify whether the TH-reduced expression in the SNpc may alter the nigrostriatal DA pathway, the potential diminished DA transmission was tested in regard to the afferent TH-expressing fibers and the co-occurrence of D1R expression with the phospho-DARPP-32 (P-DARPP-32) labeling in the striatum (Fig 4). ENZ, but not AAP, led to the inhibition of the staining intensity of the TH^+^-fibers when compared with respective vehicles (Fig 4A and B). To assess a potential impact on the DA synatic transmission, we quantified the labeling of P-DARPP-32 particularly in neurons expressing D1R likely indicative of striatal medium spiny neurons (MSNs) activation. ENZ when compared with Veh-ENZ, drastically and significantly led to inhibition of striatal P-DARPP-32 staining mostly found in NeuN^+^-neurons while P-DARPP-32 was not modified between AAP- and Veh-AAP-group conditions (Fig 4 and S2 Fig). These results suggest that ENZ is associated with a diminution of TH-expression in AR^+^-neurons in SNpc and striatal DA activity, potentially associated with ENZ-evoked reduced motivation for movement.

**Fig 4.**
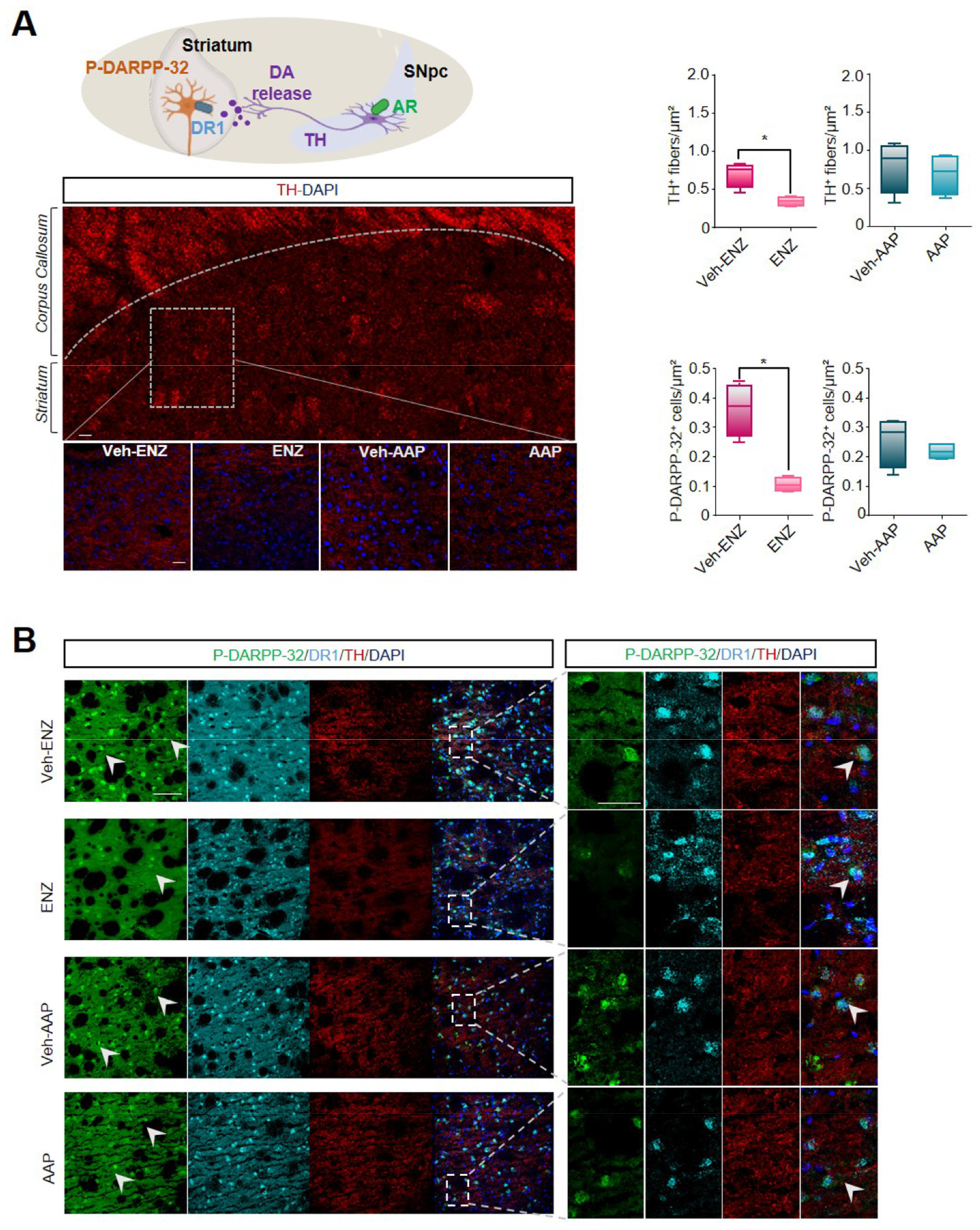
ENZ modulates midbrain dopaminergic release to striatum. (A) Schematic of SNpC dopaminergic projections to striatum and consequent striatal neuronal activation. Immunofluorescence of TH^+^-DA striatal projections of ENZ-, AAP- and vehicle-treated mice (seen on magnification of squared areas). Scale bar: 100 μm. On the right, Whisker boxes represent TH^+^ fibers intensity and the number of P-DARPP-32^+^ cells in the striatum of ENZ- and AAP-treated mice compared with respective vehicles. Statistical quantifications were performed using Mann-Whitney test. Data are represented as box and whiskers of mean□±□SEM (*n*□=□4 mice), \**p*□<□0.05, \*\**P*□≤□0.01. Scale bar: 100 μm. (B) Representative images of Dopamine cAMP-Regulated Neuronal Phosphoprotein P-DARPP-32^+^-(green), DA receptor 1^+^-(DR1, cyan), TH^+^-(red) positive fibers and DAPI (blue) immunoreactivities in striatum of ENZ-, AAP- and Vehicle-treated mice. Scale bar: 25 μm and 100 μm. AAP, abiraterone acetate-prednisone; DA, dopamine; DAPI, 4′,6-diamidino-2-phenylindol; ENZ, enzalutamide; P-DARPP-32, Phosphorylated form of Dopamine cAMP-Regulated Neuronal Phosphoprotein; DR1, DA receptor 1; SNpc, substantia nigra pars compacta; TH, Tyrosine hydroxylase; vHp, ventral hippocampus.

VTA activity can be connected to the hippocampus, to the vHP in particular where some projections are dopaminergic (41). vHp can be involved in limbic functions regulating the impact of emotional experiences (42) and controlling exploration of novelty and motivated behaviors (43). Here we detected only a modest expression of AR in the DG, as well as in CA1 and CA3 of vHp more specifically in NeuN^+^-neurons and not GFAP^+^-glial cells from ENZ-, AAP- and vehicle-treated groups (S3 Fig). To examine the impact of ENZ on VTA-dopaminergic projections to vHP neurons, the expressions of TH, P-DARPP-32 and the immediate early gene (c-fos) were quantified. TH^+^-fibers showed reduced staining intensity in the ENZ compared with Veh-ENZ group while AAP failed to alter TH labelings (Fig 5A). Moreover, in most D1R^+^-neurons, a decrease in P-DARPP-32^+^-cells and P-DARPP-32^+^-neurons (NeuN^+^-cells) (S4 Fig) in the vHp area was found in ENZ-treated brain slices (Fig 5B). This was associated with a repression of cFos^+^/NeuN^+^ neuronal cells in the vHP DG, CA3 and CA1 mainly in ENZ group (Fig 5C). These results suggest that ENZ mainly impairs vHP hippocampal activity by modulation of dopaminergic neuronal stimulation from VTA, even if the contribution of the local inhibition of AR cannot be excluded.

**Fig 5.**
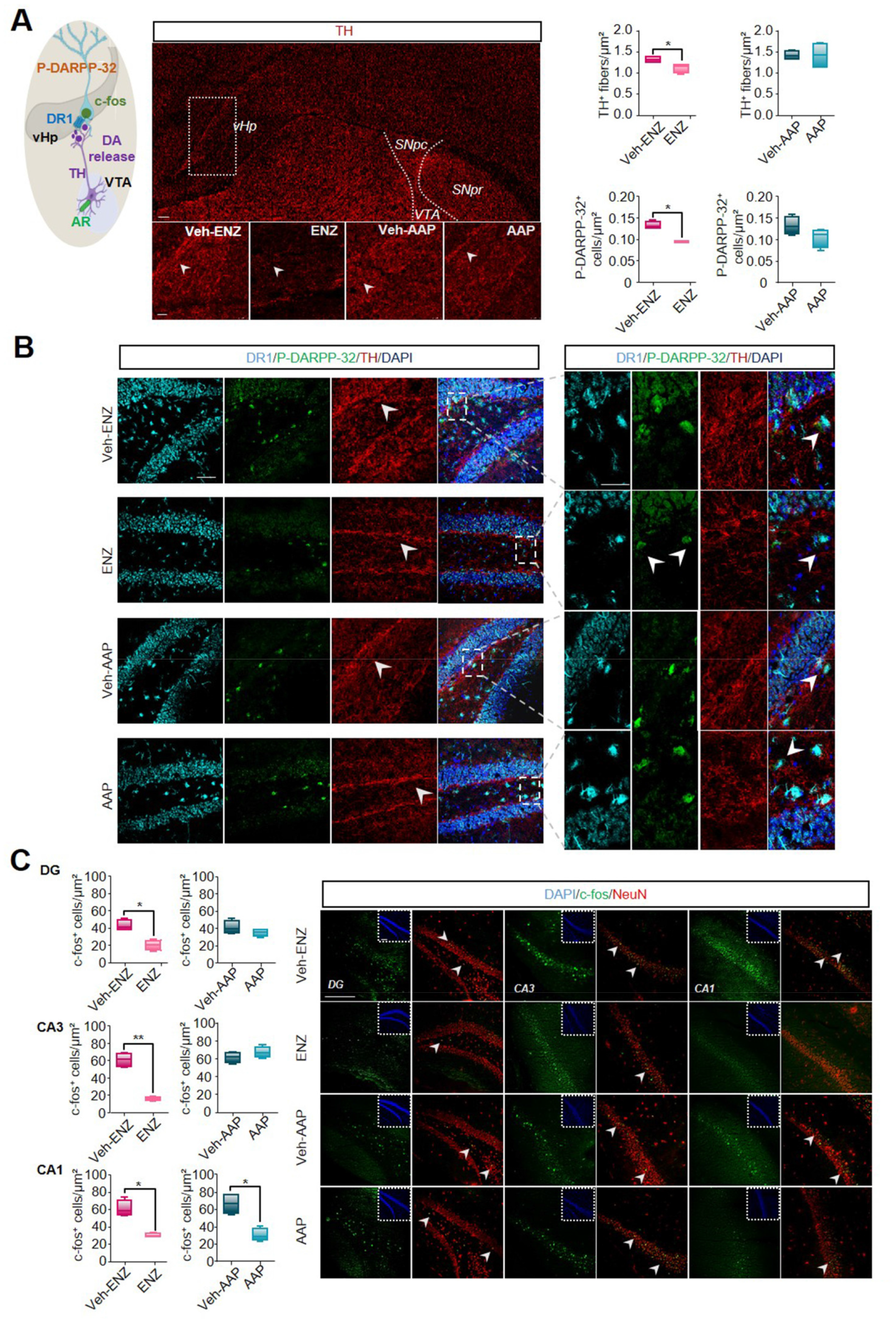
ENZ modulates midbrain dopaminergic release to ventral hippocampus. (A) Schematic representation of dopaminergic projections to ventral hippocampus (vHp) from VTA and consequent neuronal activation. Immunofluorescence of TH^+^-DA projections likely arising from VTA and projecting to vHp of ENZ-, AAP- and vehicle-treated mice (seen on magnification of squared areas). Scale bar: 100 μm. On the right, Whisker boxes represent the intensity of TH^+^ fibers and number of P-DARPP-32^+^ cells in the vHp (right panel) of ENZ- and AAP-treated mice compared with respective vehicles. Statistical quantifications were performed using Mann-Whitney test. Data are represented as box and whiskers of . Scale bar: 100 μm. (B) Representative images of P-DARPP-32^+^- (green), DR1^+^- (cyan), TH^+^- (red) positive fibers and DAPI (blue) immunoreactivities in vHp of ENZ-, AAP- and Vehicle-treated mice. Scale bar: 25 μ and 100 μm. (C) Representative example of immediate early gene (c-fos, green) and NeuN (red) immunoreactivities in the DG, CA3 and CA1 regions of vHp of ENZ-, AAP- and vehicle-treated mice. The boxed areas show DAPI (blue) immunoreactivity. On the left, wisker boxes represent the number of c-fos^+^ cells in DG, CA3 and CA1 hippocampal areas of ENZ- and AAP-treated mice compared with respective vehicles. Statistical quantifications were performed using Mann-Whitney test. Data are represented as box and whiskers and mean□±□SEM (*n*□=□4 mice), \**p*□<□05, \*\**P*□≤□0.01. Scale bar: 100 μm. AAP, abiraterone acetate-prednisone; CA1, cornus ammonis 1; CA3, cornus ammonis 3; c-fos, cellular oncogene c-fos; DA, dopamine; DAPI, 4′,6-diamidino-2-phenylindol; DG, dentate gyrus; ENZ, enzalutamide; NeuN, Neuronal Nuclei Antigen; P-DARPP-32, Phosphorylated form of Dopamine cAMP-Regulated Neuronal Phosphoprotein; DR1, DA receptor 1; TH, Tyrosine hydroxylase; vHp, ventral hippocampus; VTA, ventral tegmental area.

### ENZ treatment and not AAP specifically altered spatial learning efficiency and cognitive score

The role of androgen deprivation and direct AR antagonism on hippocampal functions and spatial working memory (44), spatial learning and memory, retrieval and behavioral flexibility was assessed by MWM Test (Fig 6A and B). No modification was detected in short-term and long-term memory using the probe and the retrieval tests respectively (Fig 6C, S3 Table). During the flexibility phase, no change was detected in ENZ- and AAP-treated groups when compared with vehicles in term of duration (Fig 6C) and distance crossed (S3 Table). During the learning phase, distance crossed, mean speed and duration were evaluated. Neither ENZ- nor AAP-treated mice exhibited significant changes in learning abilities compared with Veh (Fig 6B, S3 Table). Efficient navigation by using visual cues at disposal depends on the capacity of integration of egocentric route-based knowledge into an allocentric representation (45). We adapted a Python Neural Network to allow analysis of different swim strategies in six different classes (Fig 6D). In Veh-ENZ and Veh-AAP groups, mice used a high proportion of random swim like scanning on D1 followed by more efficient strategy on D3-4 as direct or rotating (Fig 6E). In ENZ compared with Vehicle mice, less succesful search strategies like thigmotaxis were observed (12,5 % of all strategies) contributing to a diminished cognitive score (46) only at D1 (Fig 6E) while AAP mice showed only a longer time to switch towards efficient strategies (S5 Fig).

**Fig 6.**
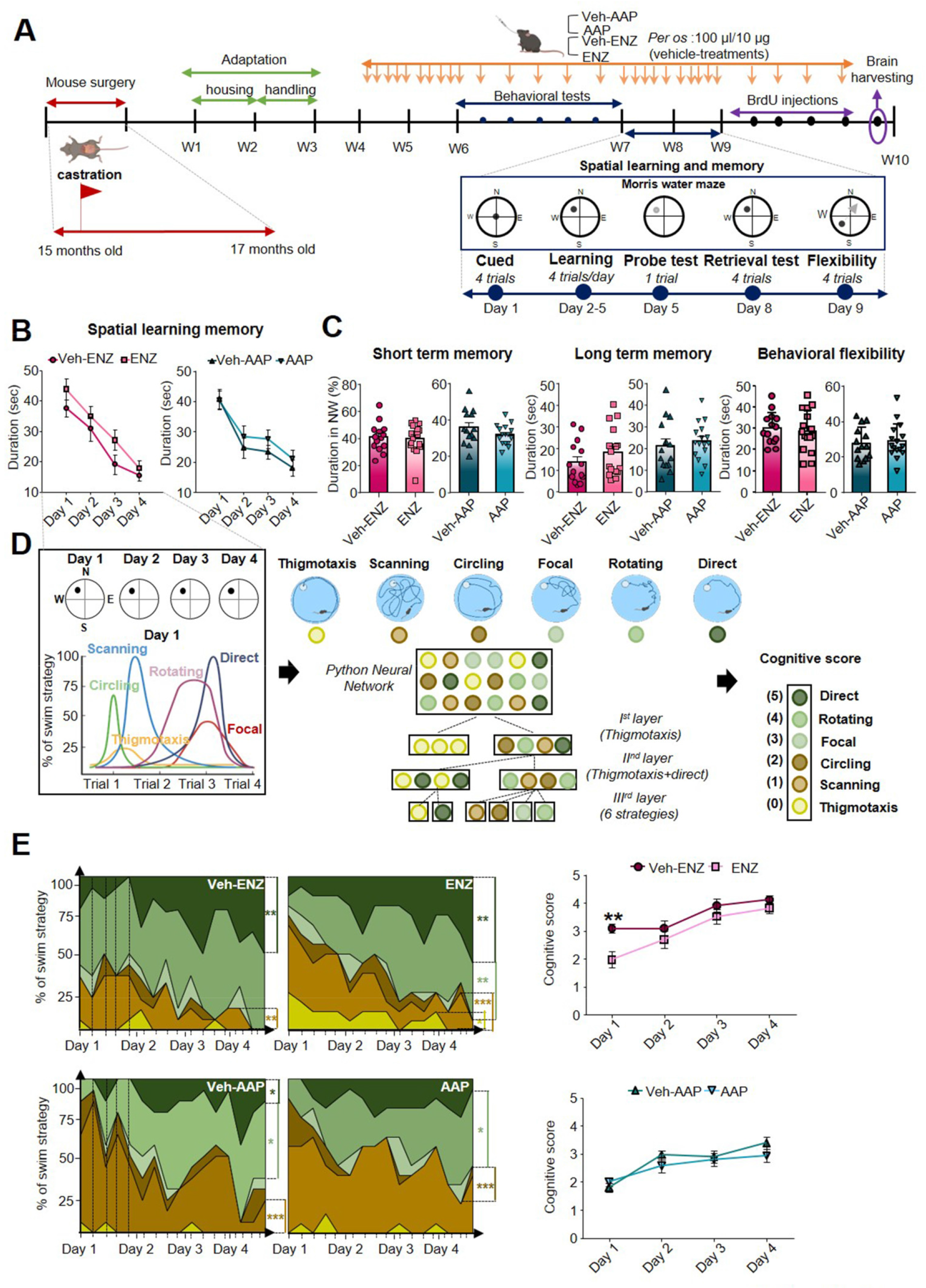
ENZ and not AAP specifically alters spatial learning efficiency and cognitive score. (A) *In vivo* experimental time course of treatment delivery and learning and memory cognitive evaluation by Morris water maze test (MWM). MWM was performed for 7 days starting from W7 (blue arrow) with familiarization (cued) on D1, the learning phase from D2 to D5, the probe test two hours after learning trials on D5, the retrieval test on D8 and the flexibility analyzed on D9 during treatments as indicated (orange arrows). Then during 4 days mice received one BrdU injection per day (50 mg/Kg) before sacrifice, brain were harvested for immunohistochemical analyses (purple arrow) and plasma were sampled. (B) Cognitive consequences of ENZ or AAP treatment compared with vehicle on spatial learning. Bar scatter plots showed the time to reach the platform (escape latency) during the learning phase. Statistical comparison of treatment groups was performed between vehicles (*n* = 14) and treated (*n* = 16) mice by repeated measures one-way ANOVA followed by Sidak’s multiple comparisons test. Bars are mean□±□EM while symbols individual data point. (C) Bar graph quantification of short-term memory (duration crossed in the target NW quadrant), long-term memory (escape latency) and behavioral flexibility (escape latency). Statistical analysis was performed between vehicle (*n* = 14) and treated (*n* = 16) mice data using unpaired *t*-test and Mann-Whitney test. (D) Left panel, schematic representation of platform positioning shown for each day of the learning phase (left panel): 4 trials with 4 different start location (N, S, W, E) were assessed each day. An illustration of swim strategy distribution along the 4 trials of day 1 of learning was drawn (left). Middle panel, visual representations of swim tracks: thigmotaxis (yellow), scanning (light brown), circling (dark brown), focal (light green), rotating (middle green) and direct (dark green). Below, simplified illustration of convolution layers of the neural network used to classify swim strategies: the starting dataset of image files corresponding to tracking trajectories in all trials along the four consecutive learning days for each mouse was run throughout the first convolution layer, which distinguishes thigmotaxis from other trajectories. The second convolution layer separates thigmotaxis and direct swim from other trajectories and the third one classifies the 6 specific trajectories. Right panel, each training trial was scored such that more efficient strategies received higher scores according to the following scale: thigmotaxis = 0, scanning = 1, circling = 2, focal = 3, rotating = 4, direct = 5. (E) Left panel, Distribution of search swim strategies during each trial for the 4 days of learning. Swim path to target are presented as stacked area plots and percentage, vehicle (*n* = 14) and treated (*n* = 16). The statistical comparison of strategy evolution from D1 to D4 was assessed by Chi-square test with Yates’ continuity correction. Right panel, statistical analysis of cognitive scores during the 4 days of learning (4 trials/day) between vehicles (*n* = 14) and treated (*n* = 16) mice by repeated measures (RM) one-way ANOVA, using the Tukey’s test for multiple comparisons. Data are expressed as aligned dot plots and mean SEM, \**p* 0.05, \*\**P* 0.01, ***P<0.001. AAP, abiraterone acetate-prednisone; BrdU, Bromodeoxyuridine; D, day; ENZ, enzalutamide; MWM, Morris water maze test; W, week.

### ENZ decreased c-fos-related activity of NeuN^+^-neurons whereas AAP interfered with the neurogenic process in the dorsal hippocampus

Some neurobiological markers including c-fos in dHP were recognized as being involved in the neural cognitive map of space (45) and in responsiveness to learning tasks in rodents (47). We observed that neither ENZ nor AAP significantly modified TH-staining in afferent fibers in dHP (Fig 7A). CYP17A1 immunostaining found in DG (Fig 7B), CA3 and CA1 (S6 Fig), more specifically in NeuN^+^, supports neurosteroidogenesis and potential local production of T or DHT in dHp (Fig 7C). The co-localization of AR as potential binding sites for locally produced androgens with BrdU^+^-neural proliferating progenitors, DCX^+^-immature hippocampal neurons or NeuN^+^-mature neurons was thus tested (Fig 7D). We did not show AR^+^/BrdU^+^ and AR^+^/DCX^+^ co-expressing cells, but observed a main proportion of AR^+^/NeuN^+^-cells in dorsal DG of dHP of ENZ-, AAP- and vehicles groups (Fig 7D). The neurogenic process and BDNF expression in dHP of ENZ-treated aged mice were not altered, but enhanced BrdU^+^- and reduced BDNF^+^-cells were measured in dHp of AAP compared with Veh-AAP mice, with no consequences on mature neurons (S7B Fig). However, exploration and overtrained responding and motor activity during spatial learning have been related to CA1 neuron activity in dHP (48). We found a significant decrease in c-fos expression in ENZ-treated mice in DG and CA1 in NeuN^+^-neurons and not in GFAP^+^-glial cells when compared with veh-ENZ (Fig 8). No difference was detected for c-fos expression in neither regions of dHP from AAP compared with Veh-AAP brain slices (Fig 8). Together, ENZ treatment is specifically associated with a decreased c-fos-related activity of DG and CA1 neurons in the dHP, that would sustain a deficit in exploratory performances.

**Fig 7.**
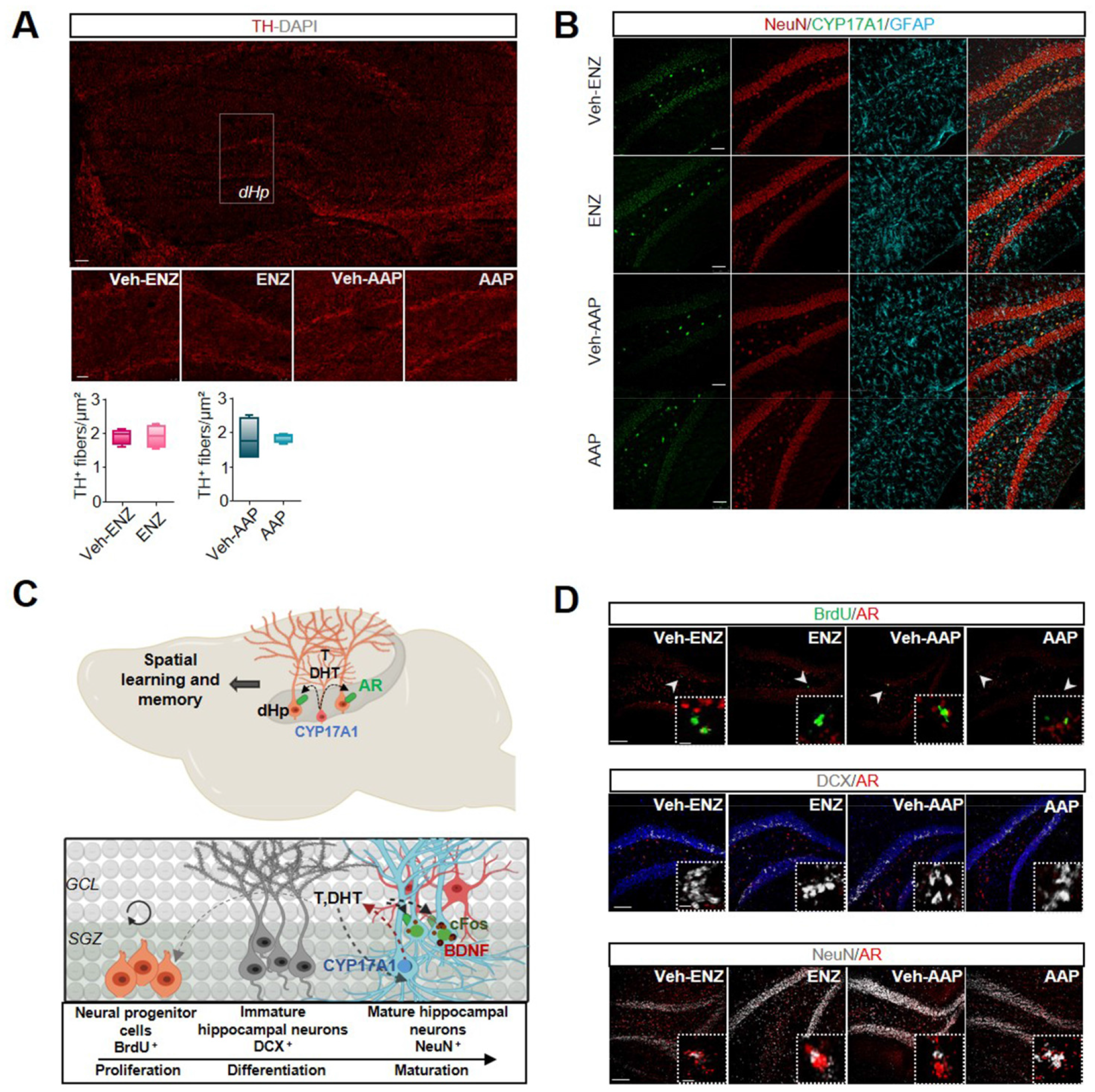
AAP and ENZ treatment impair adult neurogenesis and neuronal activity in dorsal hippocampus respectively. (A) Immunofluorescence of TH^+^-DA projections to dHP of ENZ-, AAP- and vehicle-treated mice (seen on magnification of squared areas). Whisker boxes represent the number of TH^+^ fibers in dHP. Statistical quantifications were performed using Mann-Whitney test. Bars are mean ± SEM (*n* = 4 mice). Scale bars: 100 μm. (B) Representative immunofluorescence of CYP17A1 (green), NeuN (red) and GFAP (cyan) in the DG of dHP of ENZ-, AAP- and vehicles-treated mice showing specific localization of CYP17A1 in neuronal and not glial cells. (C) Upper panel, schematic of mouse dorsal hippocampal (dHp) CYP17A1-mediating production of testosterone and DHT, potentially responsible for spatial learning deficits. Lower panel, hypothetical illustration of the impact of testosterone and DHT or prednisolone on neurogenesis. CYP17A1 expression by mature hippocampal neurons (neuron in light blue; astrocyte in red) of granule cell layer (GCL) of DG; AR is not expressed in neural progenitors cells stained by an anti-BrdU (orange) in the SGZ and by immature hippocampal neurons classically revealed by staining of an anti-DCX (grey) in the DG of vHP. AR expressed by mature hippocampal neurons revealed by anti-NeuN (light blue). AR (green) activation by T or DHT would control mature neuron (blue) activity as illustrated by c-fos immunolabeling. (D) Upper panel, representative immunolabeling of BrdU (green) and AR (red) immunofluorescence in dHP of ENZ-, AAP- and vehicles-treated mice. The boxed areas showed a magnification of lack of BrdU^+^/AR^+^ co-stained progenitor cells. Middle panel, immunolabeling of DCX (grey) and AR (red) immunofluorescence in dHp of ENZ-, AAP- and vehicle-treated mice. The boxed areas showed a magnification of lack of DCX^+^/AR^+^ co-stained immature migratory neurons in the SGZ. Lower panel, immunolabeling of NeuN (grey) and AR (red) in dHp of ENZ-, AAP- and vehicle-treated mice. The boxed areas showed a magnification of NeuN/AR co-expression in mature neurons. Scale bar: 100 μm and 50 μ AAP, abiraterone acetate-prednisone; AR, androgen receptor; BrdU, Bromodeoxyuridine; c-fos, cellular oncogene c-fos; CYP17A1, Cytochrome P450 Family 17 Subfamily A Member 1; DA, dopamine; DCX, doublecortin; DG, dentate gyrus; dHP, dorsal hippocampus; DHT, dihydrotestosterone; ENZ, enzalutamide; GCL, granule cell layer; GFAP, Glial Fibrillary Acidic Protein; NeuN, Neuronal Nuclei Antigen; SGZ, sub-granular zone; T, testosterone; TH, Tyrosine hydroxylase; vHP, ventral hippocampus.

**Fig 8.**
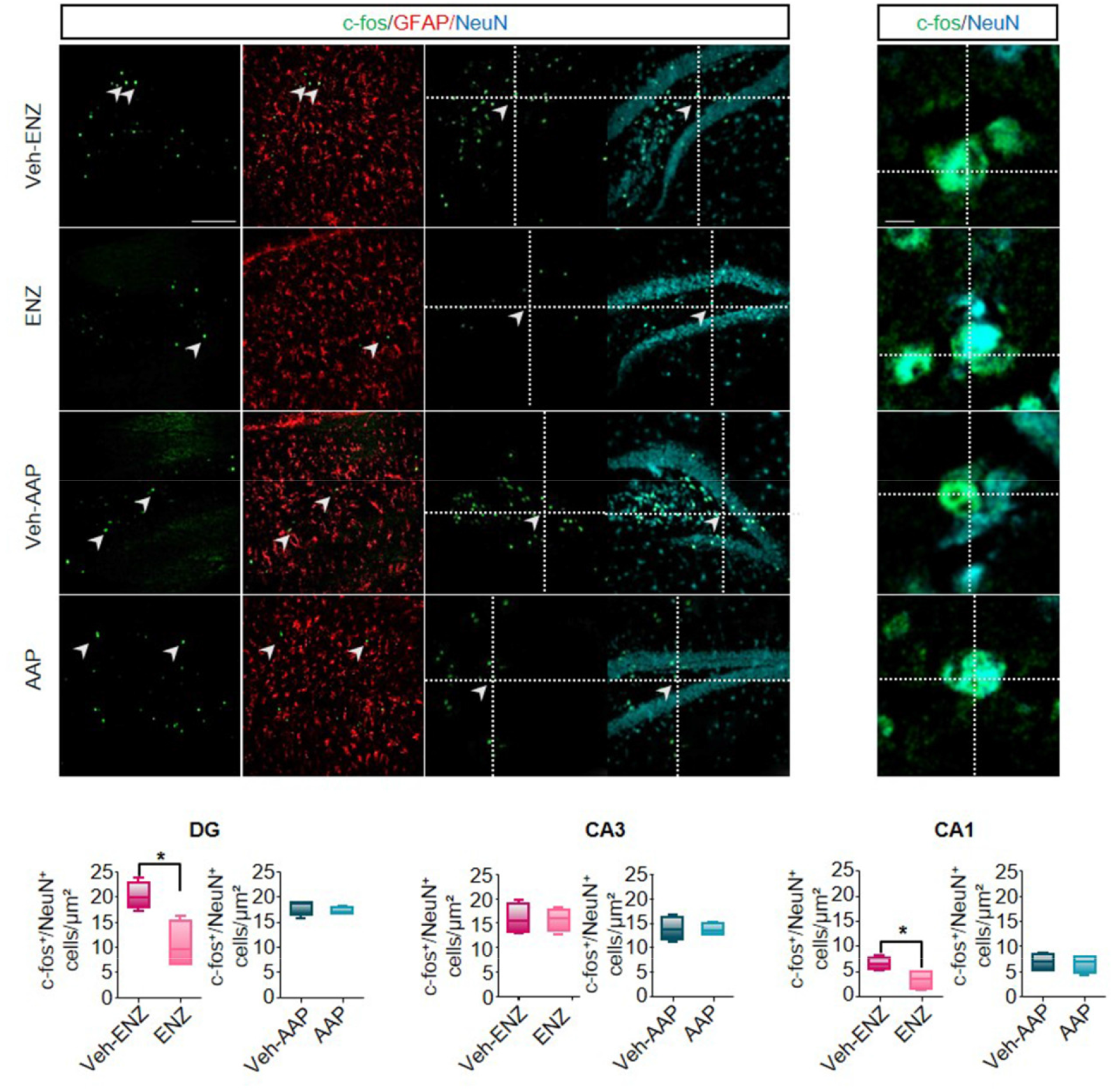
ENZ treatment impairs neuronal activity in dorsal hippocampus. Representative immunolabeling and quantification of c-fos (green) expression in NeuN^+^- (red) or GFAP^+^- (cyan) cells. Intersecting lines evidenced co-localization of NeuN^+^-cells expressing c-fos. Whisker box plots quantification of the number of c-fos^+^/NeuN^+^-cells in the DG, CA3 and CA1 areas of dHP of ENZ- and AAP-treated mice compared with respective vehicles. Statistical quantifications were performed using Mann-Whitney test. Bars are mean SEM 4 mice), \**P* < 0.05, \*\**P* 0.01. Scale bars: 100 μm. AAP, abiraterone acetate-prednisone; CA1, cornus ammonis 1 ; CA3, cornus ammonis 3; c-fos, cellular oncogene c-fos; DG, dentate gyrus; ENZ, enzalutamide; GFAP, Glial Fibrillary Acidic Protein; NeuN, Neuronal Nuclei Antigen.

## DISCUSSION

Clinical approaches to understand cognitive decline in elderly patients treated for cancer are complex and the underlying neurobiological mechanisms are far from being explained. In elderly patients with mCRPC, ENZ directly targeting AR signaling, or AAP (additioned with prednisone) inhibiting CYP17A1, significantly improved OS and QoL in two clinical trials (13, 49). However, it was shown that after 12 weeks of treatment, functional assessment of cancer therapy-prostate score improved with AAP and not with ENZ and a physical well-being subscale worsened with ENZ (24). Activation of AR signaling pathways through local synthesis of T or DHT may regulate neurotransmission, neurogenesis and brain plasticity (50), and selective behavioral responses. Here the first main evidence is that ENZ impacts locomotor and explorative behavior, and strengh capacity likely by preventing binding of central synthetized androgens to ARs expressed by DA neurons of the SNpc and the VTA. This impairs dopaminergic release in the striatum (nigrostriatal pathway) and the vHP (mesolimbocortical pathway), leading to reduced target neurons activation. The second main fact is that ENZ reduces the cognitive score indicative of learning efficiency deficit, here associated with less c-fos activation in DG and CA1 neurons of the dHP.

ADT by castration and CYP17A1 inhibition should lead to inhibition of T production and corticosterone in animals. Indeed, continuous inhibition of CYP17A1 with oral AAP in CRPC patients is safe and significantly suppresses serum androgens, estrogens and cortisol synthesis (51, 52), prednisone complementation yielding physiologic cortisol levels (53). But low levels of T can be associated with an inflammatory status in healthy elderly population (32, 33) or hypogonadism (34) while prednisone may contribute to subtle effects on immune functions (52). Inflammatory cytokines can mediate behavioral and cognitive disturbances in both human and animal subjects (32, 33), but here plasma levels of a panel of pro-inflammatory pluripotent, puripotent or chemokine cytokines were not modified in ENZ or AAP groups compared with vehicles or control mice. By discarding the possible impact of cytokines on cognitive functions (54), we thus adressed direct central mechanisms of ENZ and AAP, both penetrating the blood-brain barrier in mice (55), potentially accounting for a large part of to more fatigue and reduced activity, QoL and cognitive deficits in treated cancer elderly patients.

In aged castrated mice ENZ, and not AAP, reduced spontaneous activity and exploration, and motivation to struggle as assessed by decreased locomotor and vertical activities in OFT, increased immobility in OFT arena and TST with no difference in time immobile or latency before first immobility in FST. Immobility in TST may assess the strength and energy of the movement of mice, a sign of reluctance to maintain effort (56, 57), while as previously reported, activity in OFT can reflect neurotransmitter system effects on stimulus salience and behavioral activity (58, 59), suggesting a key role of ENZ on motivation to explore. Interestingly, pharmacological inhibition of T conversion to DHT in non castrated adolescent rats, affected exploratory and motor behaviors in OFT by down-regulating dopaminergic activity in SNpc and VTA (60). This behavioral observations were strengthened by the expression of androgen components (39) within the nigrostriatal circuitry in TH^+^-structures.

Dopaminergic pathways have been shown to drive exploratory and locomotion behaviors in rodents, promoting the positive valuation of novel stimuli as well as motivation to explore (35). In humans, dopamine has been shown to contribute to reward modulation of sensory decision-making, involving at least in part, the ventral striatum (61). Here, we show the presence of a subset of DA neurons expressing AR but also CYP17A1 in SNpc and VTA midbrain nuclei mature neurons, providing neuroanatomical bases for the local stimulation of AR by T and/or DHT throughout the nigrostriatal and mesolimbocortical pathways. There are currently few arguments in favor of a central, midbrain, expression of CYP17A1. One is that susceptibility to develop motor dysfunctions has been linked to abnormal P450 enzyme activity (62). Thus, CYP17A1 in both SNpC and VTA mature neurons may contribute to androgen synthesis and AR activation by midbrain DA neurons. This last assumption is here confirmed by the decrease of TH^+^ neuronal components expressing AR in the SNpc after ENZ treatment. Its means that dopaminergic projections from SNpc to the striatum, involved in activity or action sequence initiation and performance (63), are controlled by ENZ (Fig 9). Previous studies established that activation of DR1 and DR2-expressing neurons in the striatum relays a selective control over locomotion (64) and that P-DARPP32 can be a marker of striatal MSNs dopaminergic activation (65). Accordingly, a diminished P-DARPP32 in striatal neurons expressing at least D1R of ENZ mice, suggests that AR antagonism can lead to inhibition of striatal DA release and activity, and of motor and/or exploratory behavior. A this stage, we cannot discard that ENZ acts centrally by interacting with the GABAergic system/channel receptors (66), as previously suspected. Despite expression of CYP17A1 in SNpc and VTA, no effect of AAP is observed, potentially attributed to the weak passage across the BBB and/or a possible steroidal conversion of AA with inefficient blockage of steroidogenesis (52).

**Fig 9.**
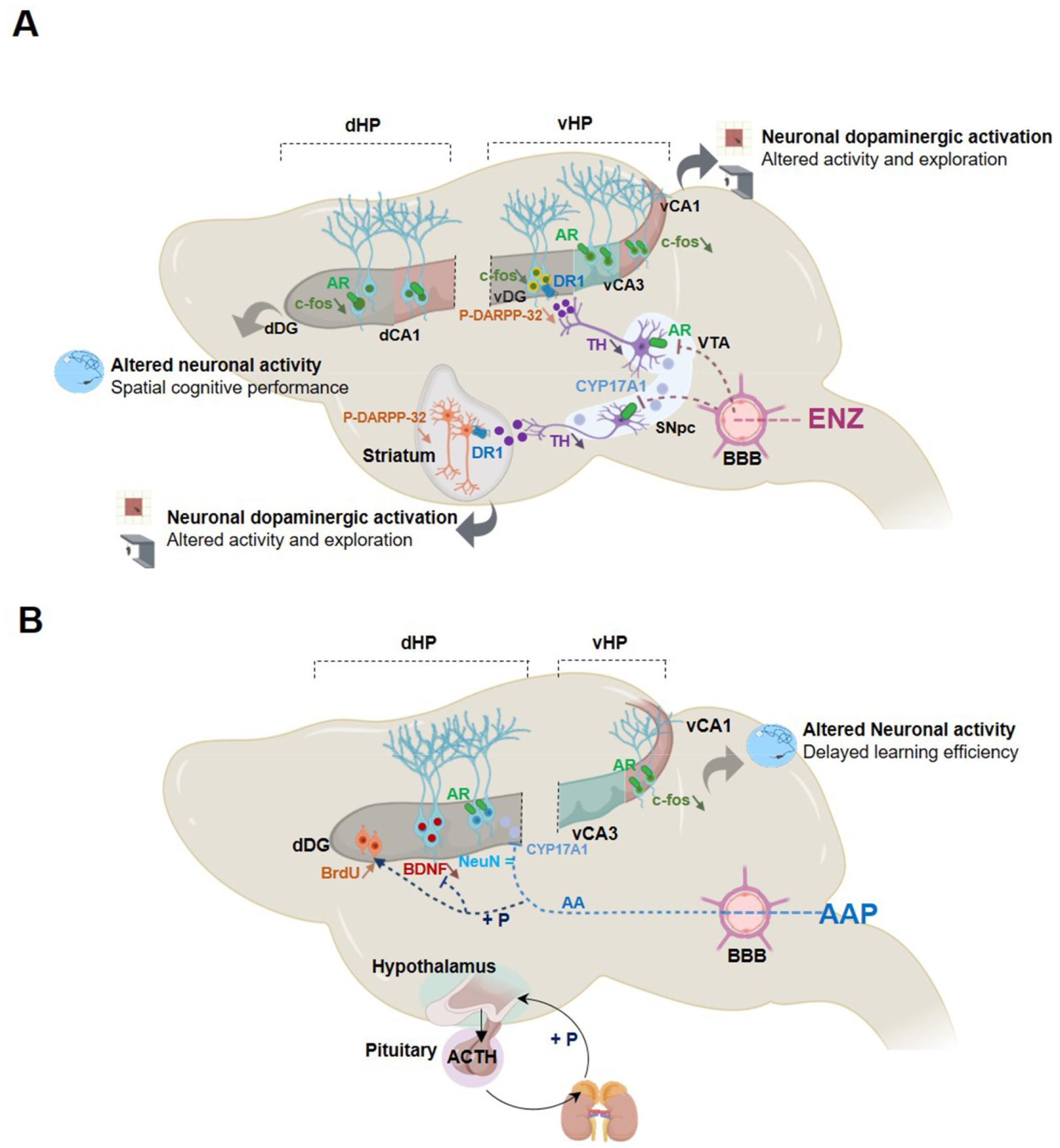
Schematic illustration of the central action of novel generation hormone therapy targeting the androgen signaling axis, abiraterone acetate+prednisone (AAP) and enzalutamide (ENZ) in aged castrated mice. (A) ENZ which is able to cross the brain blood barrier (BBB) can bind and inhibit ARs expressed in peripheral and brain organs. More specifically, CYP17A1 and AR are localized in both SNpc and VTA neurons in aged castred mouse brains. ENZ leads to reduced P-DARPP-32 activated signaling in mature neurons expressing DR1 in the striatum through the nigrostriatal pathway and in the vHP via part of the mesolimbocortical pathway. This inhibition DA-expressing afferents likely accounts for diminished locomotor activity and exploration behavior. ENZ also depressed mature neuron activity evidenced by c-fos expression in dHP potentially responsible for an altered spatial cognitive performance. (B) AAP is a prodrug, generating abiraterone selective for CYP17A1 enzyme activity, thus inhibiting androgen biosynthesis in the testis and the adrenal gland, but potentially in the VTA and SNpc. AA may have the ability to cross the BBB. In order to avoid cortisol deficiency, the treatment is combined with chronic use of prednisone (P) sufficient to activate the HPA-related feedback loop and to reduce the plasma ACTH level allowing physiological cortisol production in patients. CYP17A1 and AR are expressed in the dHP, and here the AAP effect on proliferation of precursor cells and inhibition of BDNF production without modification of mature neurons would hypothetically involve glucocorticoids receptors. In the vHP, the AAP-evoked reduction of c-Fos in mature neurons can sustain the delay in efficiency in the spatial learning acquisition. AA, abiraterone acetate; AAP, abiraterone acetate-prednisone; ACTH, adrenocorticotropic hormone; AR, androgen receptor; BBB, blood brain barrier; BDNF, Brain-derived neurotrophic factor; c-fos, cellular oncogene c-fos; CYP17A1, Cytochrome P450 Family 17 Subfamily A Member 1; DA, dopamine; dHP, dorsal hippocampus; ENZ, enzalutamide; HPA, hypothalamic–pituitary– adrenal; P, prednisone; P-DARPP-32, Phosphorylated form of Dopamine cAMP-Regulated Neuronal Phosphoprotein; DR1, DA receptor 1; SNpc, substantia nigra pars compacta; vHP, ventral hippocampus; VTA, ventral tegmental area.

DA transmission from VTA to the hippocampus has been associated with novelty detection and motivation to explore (37, 38) suggesting that the observed ENZ-induced less motivation to explore, also involved the mesolimbocortical pathway. Less TH^+^-neurons expressing AR in the VTA and TH^+^-positive fibers in the vHp of ENZ mice indicate mesolimbocortical circuit involvement. This was evidenced by a reduced DARPP32 phosphorylation and c-fos expression through the DG, CA1 and CA3, as key markers of DA neuron activity (67). In addition, AR is also found throughout the vHp supporting sensitivity of DG, CA3, and mostly CA1 to local steroidogenesis (68). As in the study showing that reduced neuronal activity in the vHP constitutes a neurobiological substrate of reduced exploratory behaviors (42), the ENZ-evoked inhibition of vHP neuron activity should contribute to depress motivation to explore and/or to effort of aged castrated mice.

Central role of androgens in spatial learning and memory abilities have been described (69–71) but in MWM test, no macroscopic impairment was detected in learning abilities, short- and long-term memories and behavioral flexibility. However, a detailed classification of swimming paths by dividing them into different “exploration strategies” (72), yielding a score based on animal relevance to spatial learning (46), highlighted that ENZ exhibit a lower cognitive score than Veh-ENZ mice. It results in prolonged occurrence of ineffective thigmotaxis behavior search strategy resulting in reduced efficacy in spatial exploration. AAP did impact cognitive score when compared with veh-AAP, showing only a slower acquisition of more effective strategies.

The hippocampus is considered “the neural substrate of a cognitive map of space” (47). dHP in particular is required for normal acquisition of a spatial memory task and learning (73), and hippocampal adult neurogenesis is necessary for a proper hippocampus-dependent spatial learning (74, 75). Stimulating effects of T or DHT on SVZ progenitor proliferation (76–78) and on newborn neuron survival (79), can be related to CYP17A1 expression in rodent hippocampus (80), mainly in CA1-CA3 pyramidal neurons (81). In our aged castrated mouse model, CYP17A1 appears sparsely but specifically expressed in mature dHP neurons In dHP, we also found AR expressed rather on dHp mature neurons instead of neural proliferating cells or immature hippocampal neurons, likely insensible to androgens. Accordingly, neither ENZ nor AAP modifies vHp neurogenesis and the number of mature neurons was similarly unmodified in dHP after treatments. Even if one observes that AAP treatment increased the number of proliferating neural cells and decreased hippocampal expression of BDNF, the lack of AR expression in these cells suggests contribution of the exogenously co-administered prednisone. Consistently the hippocampus highly express glucocorticoid receptors (82), and exposure to low concentration of glucocorticoid enhances proliferation of progenitor cells and suppresses differentiation into neurons, through decreased BDNF (82, 83). AAP-induced reduction in BDNF expression may support the slower improvement of ongoing swimming strategies in MWM, BDNF facilitating the short-term plasticity necessary for rapid switches in swim paths (84). To tentatively explain the ENZ-induced reduced performances in learning strategies during the MWM, we tested whether the VTA is the presumed source of DA in the dHP (85). Notably, DA fibers originating from VTA and contacting hippocampus control place location and reward during spatial learning (86) while VTA disruption has been shown to impair spatial performances in MWM test (87). However, the DA transmission (TH^+^-fibers) is not altered in the dHP of ENZ-treated mice, thus discarding this potential mechanism. As a marker of dHP neuron activity, we tested c-fos expression, which appears markedly reduced in dHP of ENZ-treated mice. Interestingly, inhibition of c-fos expression in dCA3 has been shown to slow down the learning process but not to prevent it (88). This can indicate that ENZ acts by antagonizing androgen signaling in dHP leading to alteration of efficient scanning of the environment to achieve the goal.

In conclusion, our findings establish the deleterious impact of ENZ on exploration, as well as on spatial learning performances in aged castrated male mice. These results pave the way for future research aimed at improving the management of QoL in patients with mCRPC to preserve from fatigue and CRCI.

## Acknowledgments

This study was enabled by the possible free accessibility to the deep learning framework Chainer. We are grateful to Lorenzo Genuardi for help in the modification of the python program and systematic evaluation of the classification accuracy of learning strategies in Morris water maze test. We thank the PRIMACEN platform (Normandy Rouen University, France) for imaging equipment and Dr. Arnaud Arabo and Mrs Julie Maucotel, for animal housing and care and for access to the behavioral equipments of the Biological Resources Department (Normandie Rouen University, France).

