## Supplementary material for "The prostate cancer therapy enzalutamide compared with abiraterone acetate/prednisone impacts motivation for exploration, spatial learning and alters dopaminergic transmission in aged castrated mice": Supp info

### Supplementary Tables

| **Proinflammatory cytokines** | Veh-ENZ | ENZ | Veh-AAPI | AAP | Control | P value |
|---|---|---|---|---|---|---|
| IL-1α | 18.9 ± 2.9 | 23.9 ± 7.7 | 23.1 ± 6.2 | 19.7 ± 3.5 | 13.9 ± 2.7 | ns. |
| IL-1β | 19.9 ± 10.2 | 9.1 ± 1.97 | 18.1 ± 5.7 | 7.5 ± 0.78 | 14.2 ± 4.7 | ns. |
| TNFα | 3.18 ± 2.4 | 0.9 ± 0.3 | 3.4 ± 1.8 | 0.4 ± 0.4 | 0.6 ± 0.4 | ns. |
| IL-12p70 | 13.1 ± 2.8 | 10.7 ±1.9 | 19 ± 6.3 | 16.9 ± 5.2 | 11.8 ± 3.1 | ns. |
| **Pluripotent cytokines** | | | | | | |
| IL-2 | 13.2 ± 1.9 | 18.9 ± 5.1 | 22.1 ± 7.8 | 26.2± 13.3 | 15.9 ± 3.7 | ns. |
| IL-4 | 8.6 ± 1.3 | 6.99 ± 1.9 | 11.4 ± 3.8 | 8.4 ± 1.9 | 8.6 ± 2.8 | ns. |
| IL-6 | 8.5 ± 1.1 | 23.4 ± 10.8 | 11.2 ± 3.7 | 8.9 ± 2.6 | 11.5 ± 3.3 | ns. |
| IL-17 | 8.8 ± 3.5 | 10.5 ± 4.7 | 14.7 ±6.7 | 0.4 ± 13.1 | 12.3 ± 3.5 | ns. |
| **Chemotactic cytokines** | | | | | | |
| RANTES | 15.9 ± 1.6 | 19.3 ± 2.1 | 16.9 ± 5.1 | 12.8 ± 3.2 | 14.3 ± 2.5 | ns. |
| MIP-1α | 2.7 ± 0.0 | 2.9 ± 0.3 | 3.1 ± 0.4 | 2.7 ± 0.0 | 5.9 ± 2.7 | ns. |
| MCP-1 | 51.6 ± 9.9 | 74.0 ± 11.2 | 59.2 ±12.3 | 40.8 ± 21.9 | 39.7 ± 11.7 | ns. |
| **Anti-inflammatory cytokines** | | | | | | |
| IL-3 | 5.5 ± 1.5 | 5.4 ± 3.8 | 4.9 ± 2.1 | 9.5 ± 5.8 | 5.2 ± 3.5 | ns. |
| **Leukocyte growth cytokines** | | | | | | |
| IL-10 | 9.8 ± 2.8 | 6.1 ± 1.9 | 4.7 ±2.4 | 13.1 ± 5.6 | 2.9 ± 1.0 | ns. |
| GM-CSF | 10.5 ± 2.0 | 10.9 ± 2.9 | 14.2 ±3.2 | 12.9 ± 5.5 | 8.1 ± 1.3 | ns. |

**S1 Table. Lack of effect of ENZ or AAP on plasma pro-inflammatory, pluripotent, anti-inflammatory, leukocyte growth and chemotactic cytokines.** Circulating cytokines were measured from plasma of Veh-ENZ-, ENZ-, Veh-AAP-, AAP-treated aged castrated (n=8) and aged non treated non castrated control mice (n=6). All data are expressed in pg/ml. Not detectable values are expressed as the half of the minimal quantity detected by the kit within this experimental sequence. Statistical quantification among the five groups was assessed using Kruskall Wallis test followed by Dunn's multiple comparison test. Data are expressed as mean ± SEM. GM-CSF Granulocyte Macrophage Colony-stimulating factor; IL Interleukine, MCP-1 Monocyte Chemotactic Protein-1; TNF-α Tumor Necrosis Factor alpha; RANTES Regulated on Activation Normal T cell expressed and secreted; MIP-1α :

Macrophage Inflammatory Protein 1-Alpha; MCP-1, Monocyte Chemotactic Protein ; GM-CSF, Granulocyte-Macrophage Colony Stimulating Factor.

| Non-treated non-castrated aged mice<br>Spontaneous activity and exploratory behaviors | Variable | Mean ± SEM |
|---|---|---|
| ***Open field test*** | | |
| Control | Vertical activity | 46.7 ± 2.499 |
| | Total distance | 30.0 ± 1.541 |
| | Total time immobile | 200.6 ± 8.835 |
| | Immobile episodes | 53.67 ± 3.403 |
| | Grooming time | 24.9 ± 4.751 |
| | Center entries | 48.33 ± 3.412 |
| | Time in center | 116.4 ± 20.43 |
| | Distance in center | 8.457 ± 0.9453 |
| | Time immobile in center | 36.12 ± 11.62 |
| | Periphery entries | 49.17 ± 3.544 |
| | Time in periphery | 483.6 ± 20.43 |
| | Distance in periphery | 21.55 ± 1.237 |
| | Time immobile in periphery | 164.5 ± 16.60 |
| **Anxiety-like behaviors** | | |
| ***Elevated plus maze*** | | |
| Control | Distance crossed | 3.816 ± 0.5591 |
| | Time immobile | 215.8 ± 14.97 |
| | Immobile episodes | 38.67 ± 2.753 |
| | SAP | 4.833 ± 1.249 |
| | Head dips | 18.67 ± 3.051 |
| | % of time in open arms | 24.00 ± 12.76 |
| | % of distance crossed in open arms | 23.17 ± 9.382 |
| ***Light dark box*** | | |
| Control | Entries in the light box | 7.167 ± 1.887 |
| | Time in the light box | 58.25 ± 18.36 |
| | Latency to the first entry in the light box | 193.3 ± 65.71 |
| **Depressive-like behaviors** | | |
| ***Tail suspension test*** | | |
| Control | Immobility duration | 175.1 ± 175.1 |
| | Latency to the first immobile episode | 78.82 ± 5.497 |
| ***Forced swim test*** | | |
| Control | Immobility duration | 163.5 ± 15.05 |

| | | |
|---|---|---|
| | Latency to the first immobile episode | 169.9 ± 15.28 |

**S2 Table. Summary of the behavioral phenotypes of non-treated non-castred aged mice.** Different items were analyzed in the open field test, the elevated plus maze test, the light and dark box test, the tail suspension test, the forced swim test in aged non castred non treated control mice (n=6). Data are expressed as mean ± SEM.

| NGT-treated aged castrated mice<br>Spontaneous activity and exploratory behaviors | Variable | Mean±SEM | $T_{DFn}$ or $F_{DFn,DFd}$ or $U_{DFn}$ | P value |
|---|---|---|---|---|
| ***Open field test*** | | | | |
| Veh-AAP vs. AAP | Vertical activity | 34.86 ± 3.477 vs. 29.94 ± 2.586 | $t_{28}$= 1.153 | P=0.2586 |
| | Total distance | 22.23 ± 1.367 vs. 20.71 ± 1.367 | $t_{28}$= 0.7839 | P=0.4397 |
| | Total time immobile | 269.9 ± 15.67 vs. 306.5 ±17.26 | $t_{28}$= 1.554 | P=0.1314 |
| | Immobile episodes | 60.79 ± 2.557 vs. 59.81 ± 2.370 | $t_{28}$= 0.2793 | P=0.7821 |
| | Grooming time | 19.49 ± 2.132 vs. 12.73 ± 1.229 | $t_{28}$= 2.835 | **P=0.0084** |
| | Center entries | 41.50 ± 3.278 vs. 36.19 ± 3.168 | $t_{28}$= 1.163 | P=0.2547 |
| | Time in center | 133.5 ± 10.30 vs. 135.5 ± 14.44 | $U_{28}$= 104.5 | P=0.7673 |
| | Distance in center | 7.644 ± 0.6105 vs. 6.658 ± 0.6097 | $t_{28}$= 1.138 | P=0.2646 |
| | Time immobile in center | 54.19 ± 9.125 vs. 68.16 ± 11.57 | $U_{28}$= 92 | P=0.4232 |
| | Periphery entries | 41.29 ± 3.304 vs. 36.31 ± 3.162 | $t_{28}$= 1.086 | P=0.2868 |
| | Time in periphery | 466.5 ± 10.30 vs. 464.5 ± 14.44 | $U_{28}$= 104.5 | P=0.7673 |
| | Distance in periphery | 14.59 ± 0.9904 vs. 14.05 ± 0.9962 | $t_{28}$= 0.3729 | P=0.7073 |
| | Time immobile in periphery | 215.6 ± 12.78 vs. 244.6 ± 18.29 | $t_{28}$= 1.261 | P=0.2176 |
| Veh-ENZ vs. ENZ | Vertical activity | 49.62 ± 4.3999 vs. 34.67 ± 2.433 | $t_{28}$= 3.081 | **P=0.0048** |
| | Total distance | 19.54 ± 1.212 vs. 15.48 ± 0.8989 | $t_{28}$= 2.037 | **P=0.0468** |
| | Total time immobile | 251.9 ± 15.68 vs. 317.3 ± 14.92 | $t_{28}$= 3.016 | **P=0.0057** |
| | Immobile episodes | 56.54 ± 2.667 vs. 57.33 ± 2.341 | $t_{28}$= 0.2250 | P=0.8238 |

| | | | | |
|---|---|---|---|---|
| | Grooming time | 26.68 ± 3.771 vs. 29.51 ± 3.913 | $t_{28}$= 0.5152 | P=0.6108 |
| | Center entries | 26.69 ± 1.834 vs. 18.67 ± 2.042 | $t_{28}$= 2.2886 | **P=0.0077** |
| | Time in center | 49.41 ± 5.668 vs. 36.72 ± 4.777 | $t_{28}$= 1.724 | P=0.0965 |
| | Distance in center | 3.224 ± 0.2445 vs. 2.729 ± 0.3111 | $t_{28}$= 0.9668 | P=0.3419 |
| | Time immobile in center | 13.11 ± 4.368 vs. 8.525 ± 2.105 | $U_{28}$= 95.50 | P=0.5403 |
| | Periphery entries | 26.92 ± 1.893 vs. 18.44 ± 1.962 | $t_{28}$= 3.067 | **P=0.0049** |
| | Time in periphery | 550.6 ± 5.669 vs. 563.9 ± 4.514 | $t_{28}$= 1.368 | P=0.0731 |
| | Distance in periphery | 16.32 ± 1.018 vs. 12.47 ± 0.7010 | $t_{28}$= 3.202 | **P=0.0035** |
| | Time immobile in periphery | 237.8 ± 15.69 vs. 313 ± 15.10 | $t_{28}$= 3.430 | **P=0.0093** |
| **Anxiety-like behaviors** | | | | |
| ***Elevated plus maze*** | | | | |
| Veh-AAP vs. AAP | Distance crossed | 4.376 ± 0.3012 vs. 4.870 ± 0.3170 | $t_{28}$= 1.12 | P=0.2721 |
| | Time immobile | 211.1 ± 6.169 vs. 207 ± 6.406 | $t_{28}$= 0.4596 | P=0.6949 |
| | Immobile episodes | 35 ± 1.671 vs. 39 ± 1.886 | $t_{28}$= 1.567 | P=0.1283 |
| | SAP | 3.5 ± 0.9361 vs. 3.125 ± 0.7004 | $t_{28}$= 0.3256 | P=0.7471 |
| | Head dips | 12.64 ± 1.830 vs. 17.38 ± 1.248 | $t_{28}$= 2.182 | **P=0.0376** |
| | % of time in open arms | 17.14 ± 5.130 vs. 22.63 ± 4.904 | $U_{28}$= 89.50 | P=0.3588 |
| | % of distance crossed in open arms | 21.79 ± 4.497 vs. 21.31 ± 3.931 | $U_{28}$= 107.5 | P=0.8616 |
| Veh-ENZ vs. ENZ | Distance crossed | 4.347 ± 0.4035 vs. 3.818 ± 0.3838 | $t_{28}$= 0.9957 | P=0.3279 |
| | Time immobile | 206.8 ± 8.456 vs. 218.2 ± 7.562 | $t_{28}$= 0.901 | P=0.3753 |
| | Immobile | 38.07 ± 2.822 vs. | $t_{28}$= 0.5042 | P=0.6181 |

| | | | | |
|---|---|---|---|---|
| | episodes | 37.29 ± 2.230 | | |
| | SAP | 14 ± 1.383 vs. 13.44 ± 1.505 | $t_{28}$= 0.2723 | P=0.7874 |
| | Head dips | 14.43 ± 1.806 vs. 11.25 ± 1.442 | $t_{28}$= 0.2723 | P=0.7874 |
| | % of time in Open Arms | 24.22 ± 3.555 vs. 20.15 ± 4.310 | $U_{28}$= 85 | P=0.2751 |
| | % of distance crossed in open arms | 25.63 ± 3.176 vs. 18.38 ± 3.015 | $t_{28}$= 1653 | P=0.1095 |
| ***Light/dark box*** | | | | |
| Veh-AAP vs. AAP | Entries in the light box | 4.429 ± 0.7010 vs. 5.313 ± 0.6565 | $t_{28}$= 0.9203 | P=0.3653 |
| | Time in the light box | 26.26 ± 4.74 vs. 38.78 ± 5.884 | $t_{28}$= 1.625 | P=0.1153 |
| | Latency to the first entry in the light box | 213.3 ± 37.67 vs. 158.7 ± 25.85 | $t_{28}$= 1.209 | P=0.1794 |
| Veh-ENZ vs. ENZ | Entries in the light box | 8.286 ± 1.404 vs. 6.188 ± 0.7595 | $t_{28}$= 1.360 | P=0.4140 |
| | Time in the light box | 57.34 ± 9.305 vs. 50.44 ± 8.887 | $t_{28}$= 0.5367 | P=0.1560 |
| | Latency to the first entry in the light box | 139.6 ± 22.76 vs. 142.3 ± 32.87 | $t_{28}$= 0.00639 | P=0.9476 |
| **Depressive-like behaviors** | | | | |
| ***Tail suspension test*** | | | | |
| Veh-AAP vs. AAP | Immobility duration | 169.9 ± 11.31 vs. 163.2 ± 8.347 | $t_{28}$= 0.568 | P=0.5746 |
| | Latency to the first immobile episode | 103.8 ± 9.167 vs. 90.13 ± 8.161 | $t_{28}$= 1.268 | P=0.2125 |
| Veh-ENZ vs. ENZ | Immobility duration | 141.9 ± 7.083 vs. 161.3 ± 8.677 | $t_{28}$= 2.357 | **P=0.0256** |
| | Latency to the first immobile episode | 73.85 ± 6.016 vs. 73.53 ± 6.938 | $t_{28}$= 0.2267 | P=0.8223 |
| ***Forced swim test*** | | | | |
| Veh-AAP vs. AAP | Immobility | 201.2 ± 19.60 vs. | $t_{28}$= 0.3993 | P=0.6670 |

| | | | | |
|---|---|---|---|---|
| | duration | 190.3 ± 18.76 | | |
| | Latency to the first immobile episode | 117.2 ± 12.94 vs. 128.7 ± 12.53 | $t_{28}$= 0.5263 | P=0.6028 |
| Veh-ENZ vs. ENZ | Immobility duration | 206.3 ± 12.74 vs. 196.8 ± 14.26 | $t_{28}$= 0.857 | P=0.3987 |
| | Latency to the first immobile episode | 129.6 ± 15.88 vs. 136.9 ± 15.69 | $t_{28}$= 0.1279 | P=0.8991 |
| **Spatial learning and memory** | | | | |
| ***Morris water maze*** | | | | |
| Veh-AAP vs. AAP | Escape latency (familiarization) | 32.99 ± 3.315 vs. 35.23 ± 2.926 | $t_{28}$= 0.5087 | P=0.6150 |
| | Distance crossed (familiarization) | 5.417 ± 0.6168 vs. 6.067 ± 0.6149 | $U_{28}$= 97 | P=0.5521 |
| | Mean speed (familiarization) | 0.1654 ± 0.006 vs. 0.1720 ± 0.005 | $U_{28}$= 97.50 | P=0.5589 |
| | Escape latency (learning) | | Trt:$F_{1,28}$=1.131 | P=0.2966 |
| | Day 1 | 40.839 ± 3.226 vs. 40.344 ± 3.119 | | |
| | Day 2 | 24.586 ± 3.275 vs. 28.405 ± 3.484 | Day:$F_{3,84}$=19.16 | P<0.0001 |
| | Day 3 | 23.345 ± 2.282 vs. 27.612 ± 3.074 | | |
| | Day 4 | 18.029 ± 2.689 vs. 21.063 ± 2.891 | Int:$F_{3,84}$=0.2816 | P=0.8386 |
| | Distance crossed (learning) | | Trt:$F_{1,28}$=0.9956 | P=0.3269 |
| | Day 1 | 7.621 ± 0.565 vs. 7.545 ± 0.662 | | |
| | Day 2 | 4.284 ± 0.519 vs. 5.397 ± 0.678 | Day:$F_{3,84}$=21.38 | P<0.0001 |
| | Day 3 | 4.526 ± 0.445 vs. 5.048 ± 0.532 | | |
| | Day 4 | 3.417 ± 0.504 vs. | Int:$F_{3,84}$=0.4517 | P=0.7167 |

| | | | | |
|---|---|---|---|---|
| | | 3.857 ± 0.527 | | |
| | Mean speed (learning) | | $Trt: F_{1,28}=1.490$ | P=0.2324 |
| | Day 1 | 0.207 ± 0.014 vs. 0.207 ± 0.013 | | |
| | Day 2 | 0.214 ± 0.029 vs. 0.197 ± 0.007 | $Day: F_{3,84}=0.6293$ | P=0.5981 |
| | Day 3 | 0.240 ± 0.020 vs. 0.211 ± 0.015 | | |
| | Day 4 | 0.208 ± 0.015 vs. 0.0208 ± 0.012 | $Int: F_{3,84}=0.3313$ | P=0.8027 |
| | Duration (probe test) | 35.93 ± 2.567 vs. 31.75 ± 1.377 | $Trt: F_{1,28}=1.486$ | P=0.1485 |
| | Distance (probe test) | 33.21 ± 2.427 vs. 30.19 ± 1.330 | $Trt: F_{1,28}=1.131$ | P=0.2676 |
| | % of time spent in NW quadrant (retrieval test) | 21.19 ± 3.180 vs. 23.48 ± 2.143 | $t_{28}= 0.6118$ | P=0.5456 |
| | % of distance crossed in the NW quadrant (retrieval test) | 3.901 ± 0.5951 vs. 4.191 ± 0.3489 | $t_{28}= 0.4329$ | P=0.6684 |
| | Escape latency (flexibility) | 46.95 ± 4.715 vs. 45.11 ± 4.398 | $U_{28}= 99$ | P=0.5753 |
| | Distance crossed (flexibility) | 8.792 ± 0.7359 vs. 9.650 ± 0.9469 | $t_{28}= 0.7006$ | P=0.4894 |
| | Mean speed (flexibility) | 0.2029 ± 0.01436 vs. 0.1943 ± 0.008442 | $U_{28}= 105.5$ | P=0.7974 |
| Veh-ENZ vs. ENZ | Escape latency (familiarization) | 48.21 ± 3.233 vs. 47.76 ± 1.871 | $t_{28}= 0.1242$ | P=0.9020 |
| | Distance crossed (familiarization) | 8.263 ± 0.7647 vs. 8.279 ± 0.4464 | $t_{28}= 0.0180$ | P==0.9857 |
| | Mean speed (familiarization) | 0.1642 ± 0.009 vs. 0.1714 ± 0.006 | $t_{28}= 0.6618$ | P=0.5135 |
| | Escape latency (learning) | | $Trt: F_{1,28}=2.596$ | P=0.1183 |

| | | | |
|---|---|---|---|
| Day 1 | 37.655 ± 2.775 vs. 43.853 ± 3.462 | $t_{28}$= 0.6618 | P=0.5135 |
| Day 2 | 30.862 ± 4.004 vs. 34.938 ± 3.251 | Day:$F_{3,84}$=35.16 | P<0.0001 |
| Day 3 | 19.098 ± 3.203 vs. 27.073 ± 3.351 | | |
| Day 4 | 15.485 ± 1.924 vs. 17.847 ± 2.612 | Int:$F_{3,84}$=0.4674 | P=0. 7058 |
| Distance crossed (learning) | | Trt:$F_{1,28}$=2.131 | P=0.1555 |
| Day 1 | 6.581 ± 0.527 vs. 7.242 ± 0.589 | | |
| Day 2 | 5.185 ± 0.692 vs. 5.804 ± 0.549 | Day:$F_{3,84}$=27.65 | P<0.0001 |
| Day 3 | 3.185 ± 0.460 vs. 4.559 ± 0.612 | | |
| Day 4 | 2.772 ± 0.394 vs. 3.157 ± 0.474 | Int:$F_{3,84}$=0.4124 | P=0.7445 |
| Mean speed (learning) | | Trt:$F_{1,28}$=0.1416 | P=0.7095 |
| Day 1 | 0.176 ± 0.008 vs. 0.168 ± 0.007 | | |
| Day 2 | 0.172 ± 0.006 vs. 0.166 ± 0.010 | Day:$F_{3,84}$=51.72 | P=0.6715 |
| Day 3 | 0.172 ± 0.010 vs. 0.172 ± 0.009 | | |
| Day 4 | 0.176 ± 0.008 vs. 0.176 ± 0.007 | Int:$F_{3,84}$=0.3018 | P=0.8240 |
| % of time spent in NW quadrant (probe test) | 40.89 ± 2.830 vs. 39.82 ± 2.757 | $U_{28}$= 109 | P=0.9185 |
| % of distance crossed in the NW quadrant (probe test) | 39.79 ± 2.459 vs. 38.42 ± 2.496 | $U_{28}$= 110 | P=0.9510 |
| % of time spent | 13.75 ± 2.531 vs. | $t_{28}$= 1.206 | P=0.2378 |

| | | | |
|---|---|---|---|
| in NW quadrant (retrieval test) | 18.35 ± 2.800 | | |
| % of distance crossed in the NW quadrant (retrieval test) | 2.532 ± 0.4029 vs. 3.598 ± 0.6047 | $t_{28}$= 1.423 | P=0.1657 |
| Escape latency (flexibility) | 34.50 ± 5.496 vs. 29.33 ± 5.819 | $U_{28}$=97 | P=0.5409 |
| Distance crossed (flexibility) | 6.936 ± 1.088 vs. 4.935 ± 1.053 | $t_{28}$= 1.318 | P=0.1983 |
| Mean speed (flexibility) | 0.2032 ± 0.009 vs. 0.1797 ± 0.012 | $t_{28}$= 1.473 | P=0.1519 |

**S3 Table. Summary of the behavioral phenotypes of aged castred mice treated with ENZ or AAP.** Different items were analyzed in the open field test, the elevated plus maze test, the light and dark box test, the tail suspension test, the forced swim test and the Morris water maze in mice treated with veh-AAP (n=14), AAP (n=16), veh-ENZ (n=14), ENZ (n=16). Based on the group normality, data were analyzed using either: *(i)* one way ANOVA with repeated measures and t-test for parametric analysis or Mann-Whitney test for non parametric analysis. Behavioral data are expressed as mean ± SEM and statistical data are indicated as t or F value in case of parametric tests, while U value is indicated in case of non parametric test. In ANOVA analysis, the presented factors are treatment (Trt) and days and their interaction (Int). P<0.05 was considered as significant.

**Legends to supplementary figures**

**S1 Fig. ENZ or AAP have no impact on body weight gain.** (A) Body weight gain curves of ENZ-treated mice compared with Veh-ENZ. Statistical analysis was performed between vehicle (n=14) and treated (n=16) data using one-way ANOVA with repeated measures, followed by Sidak's multiple comparison test. Data are expressed as aligned dot plot and mean ± SEM. (B) Body weight gain curves of AAP-treated mice compared with Veh-AAP. Statistical analysis was performed between vehicle (n=14) and treated (n=16) data using one - way ANOVA with repeated measures, followed by Sidak's multiple comparison test. Data are expressed as aligned dot plot and mean ± SEM. AAP, abiraterone acetate-prednisone; ENZ, enzalutamide.

**S2 Fig. ENZ but not AAP decreases dopaminergic activation of striatal medium spiny neurons.** Schematic of SNpC dopaminergic projections to striatum and consequent striatal neuronal activation. Below, representative images and quantification of cAMP-Regulated Neuronal Phosphoprotein (P-DARPP-32, green) and Neuronal Nuclei Antigen (NeuN, red) immunoreactivities in brain striatum of ENZ-, AAP- and Vehicle-treated mice. Box and Whiskers (right panel) represent the number of P-DARPP-32$^{+}$/ NeuN$^{+}$ cells in the striatal area of ENZ- and AAP-treated mice compared with respective vehicles. Statistical quantification was performed by using Mann-Whitney test. Data are represented as box and whiskers and mean ± SEM (n = 4 mice), $*P < 0.05$. Scale bar: 250 μm. AAP, abiraterone acetate-prednisone; ENZ, enzalutamide; NeuN, Neuronal Nuclei Antigen; P-DARPP-32, Phosphorylated form of Dopamine cAMP-Regulated Neuronal Phosphoprotein.

**S3 Fig. Androgen receptors are expressed by mature neurons of ventral hippocampus.** Schematic (upper panel) and representative images (lower panel) of androgen receptors (AR, green), Neuronal Nuclei Antigen (NeuN, purple) and Glial Fibrillary Acidic Protein (GFAP, red) immunoreactivities in ventral DG (vDG), CA3 (vCA3) and CA1 (vCA1) of ENZ-, AAP- and Vehicle-treated mice. The boxed areas show a magnification of $NeuN^+/AR^+$ and the lack of $GFAP^+/AR^+$ cells. Scale bar: 100 μm and 50 μm . AAP, abiraterone acetate-prednisone; AR, androgen receptor; CA1, Cornu ammonis 1; CA3, Cornu ammonis 3; ENZ, enzalutamide; GFAP, Glial Fibrillary Acidic Protein; NeuN, Neuronal Nuclei Antigen; vDG, ventral dentate gyrus.

**S4 Fig. ENZ but not AAP decreases dopaminergic activation of mature neurons of ventral hippocampus.** Schematic representation of VTA dopaminergic projections to vHP and consequent neuronal activation. Below, representative images and quantification of NeuN (green) and P-DARPP-32 (red) immunoreactivities in the vDG of ENZ-, AAP- and vehicle-treated mice. A magnification of the squared area shows $P\text{-}DARPP\text{-}32^+/NeuN^+$ neurons of pyramidal tract of vHp. Box and Whiskers (right panel) represent the number $P\text{-}DARPP\text{-}32^+/$ $NeuN^+$ cells in the vDG of ENZ- and AAP-treated mice when compared with respective vehicles. Statistical quantification was performed by using Mann-Whitney test. Data are represented as box and whiskers and mean ± SEM (n = 4 mice), $*P < 0.05$. Scale bar: 100 μm.and 25 μm. AAP, abiraterone acetate-prednisone; ENZ, enzalutamide; NeuN, Neuronal Nuclei Antigen; P-DARPP-32, Phosphorylated form of Dopamine cAMP-Regulated Neuronal Phosphoprotein; vDG, ventral dentate gyrus; vHP, ventral hippocampus; VTA, ventral tegmental area.

**S5 Fig. ENZ or AAP treatment impacts on learning strategy trend overtime.** Distribution of search-strategies during the 4 days of learning. Statistical comparison of each swim strategy percentage between vehicle ($n$ = 14) and treated ($n$ = 16) mice by Chi-square test with Yates' continuity correction was assessed, each day of learning, from D1 to D4 of ENZ-

and AAP-treated mice compared with respective vehicles. Swim paths are presented as histogram plot and percentage, $*P < 0.05$, $**P \leq 0.01$. AAP, abiraterone acetate-prednisone; ENZ, enzalutamide.

**S6 Fig. CYP17A1 expression in CA3 and CA1 of dorsal hippocampus.** (A) Schematic representation of CYP17A1 expression in mature neurons of dCA3. Below, representative example of CYP17A1 (green), NeuN (red) and DAPI (blue) immunoreactivities in the dCA3 of dHP of ENZ-, AAP- and vehicle-treated mice. Intersection of horizontal and vertical lines indicates CYP17A1$^+$/NeuN$^+$ cells in the 4 conditions of treatment. The boxed areas show a magnification of CYP17A1$^+$/NeuN$^+$ cells. Scale bar: 100 μm and 50 μm. (B) Schematic representation of CYP17A1 expression in mature neurons of dCA1. Below, representative example of CYP17A1 (green), NeuN (red) and DAPI (blue) immunoreactivities in the dCA1 of dHP of ENZ-, AAP- and vehicle-treated mice. Intersection of horizontal and vertical lines indicates CYP17A1$^+$/NeuN$^+$ cells in the 4 conditions of treatment. The boxed areas show a magnification of CYP17A1$^+$/NeuN$^+$ cells. Scale bar: 100 μm and 50 μm. AAP, abiraterone acetate-prednisone; CA1, cornu ammonis 1; CYP17A1, Cytochrome P450 Family 17 Subfamily A Member 1; DAPI, 4′,6-diamidino-2-phenylindol; dCA3, dorsal Cornu Ammonis 3; ENZ, enzalutamide; HP, hippocampus; NeuN, Neuronal Nuclei Antigen.

**S7 Fig. AAP treatment alters neurogenesis with no major effect on mature neurons.** (A) Representative images and quantification of NeuN (green) and BrdU (red) immunoreactivities in the dorsal (upper panel) and ventral (lower panel) DG of ENZ-, AAP- and vehicle-treated mice. White arrows and the boxed areas show BrdU$^+$-cells and a magnification, respectively. Box and whiskers represent the number of BrdU$^+$ and NeuN$^+$ cells in dorsal (left panel) and ventral (right panel) DG of ENZ and AAP treated mice when compared with respective vehicles. Statistical quantification was performed using Mann-Whitney test. Bars are mean ± SEM (n = 4 mice), *$P < 0.05$. Scale bar: 100 μm. (B) Representative immuno-labeling of BDNF (red) (cell nuclei stained with DAPI, blue) and quantification in the dDG of ENZ, AAP and vehicle-treated mice. Statistical quantification was performed using Mann-Whitney test. Bars are mean ± SEM (n = 4 mice), *$P < 0.05$. Scale bars: 100 μm. AAP, abiraterone acetate-prednisone; BDNF, Brain-derived neurotrophic factor; BrdU, Bromodeoxyuridine; DAPI, 4′,6-diamidino-2-phenylindol; dDG, dorsal dentate gyrus; ENZ, enzalutamide; NeuN, Neuronal Nuclei Antigen.
