## Supplementary material for "The prostate cancer therapy enzalutamide compared with abiraterone acetate/prednisone impacts motivation for exploration, spatial learning and alters dopaminergic transmission in aged castrated mice": Supp figures

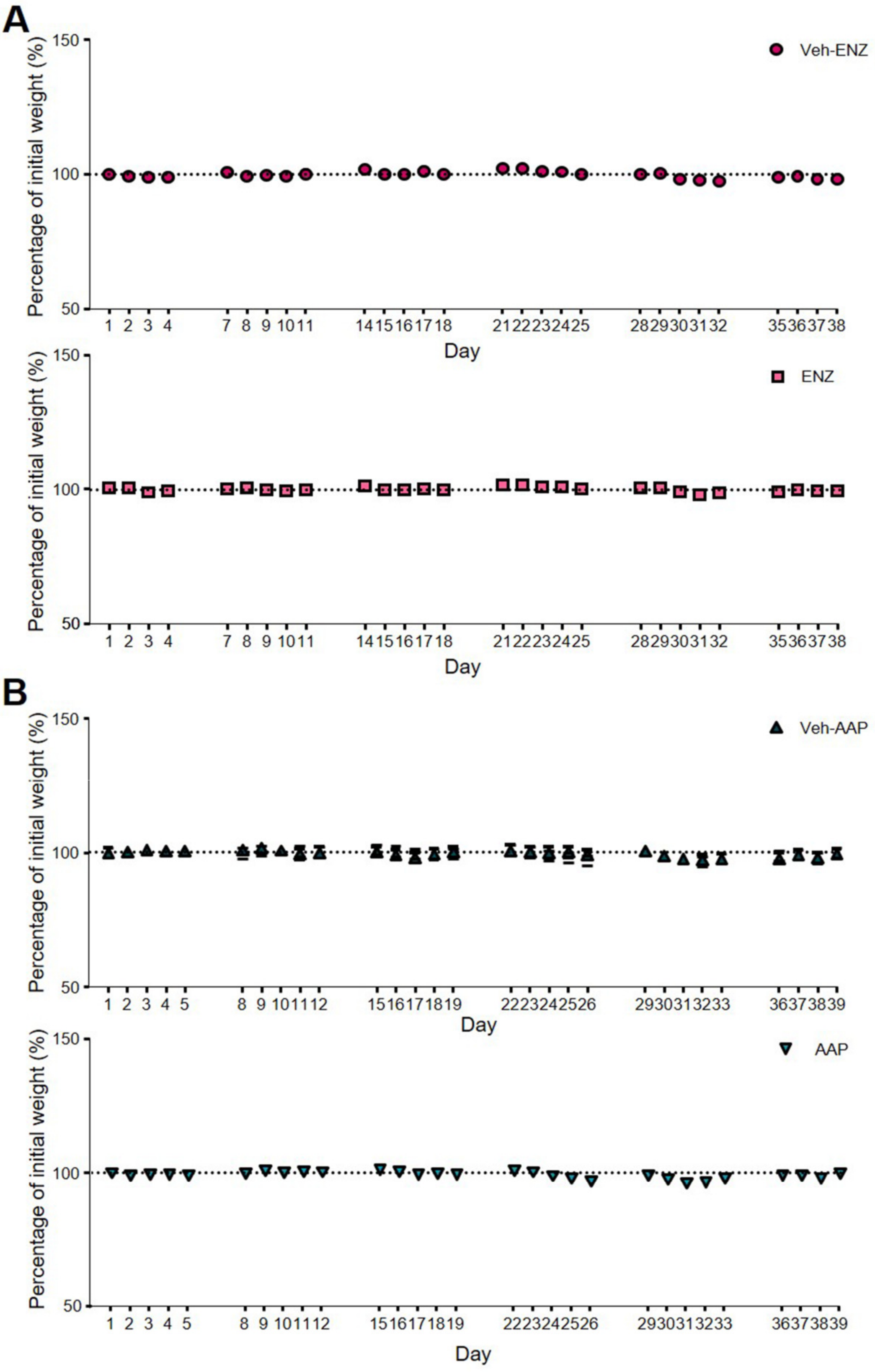


Nicola et al., S1 Fig

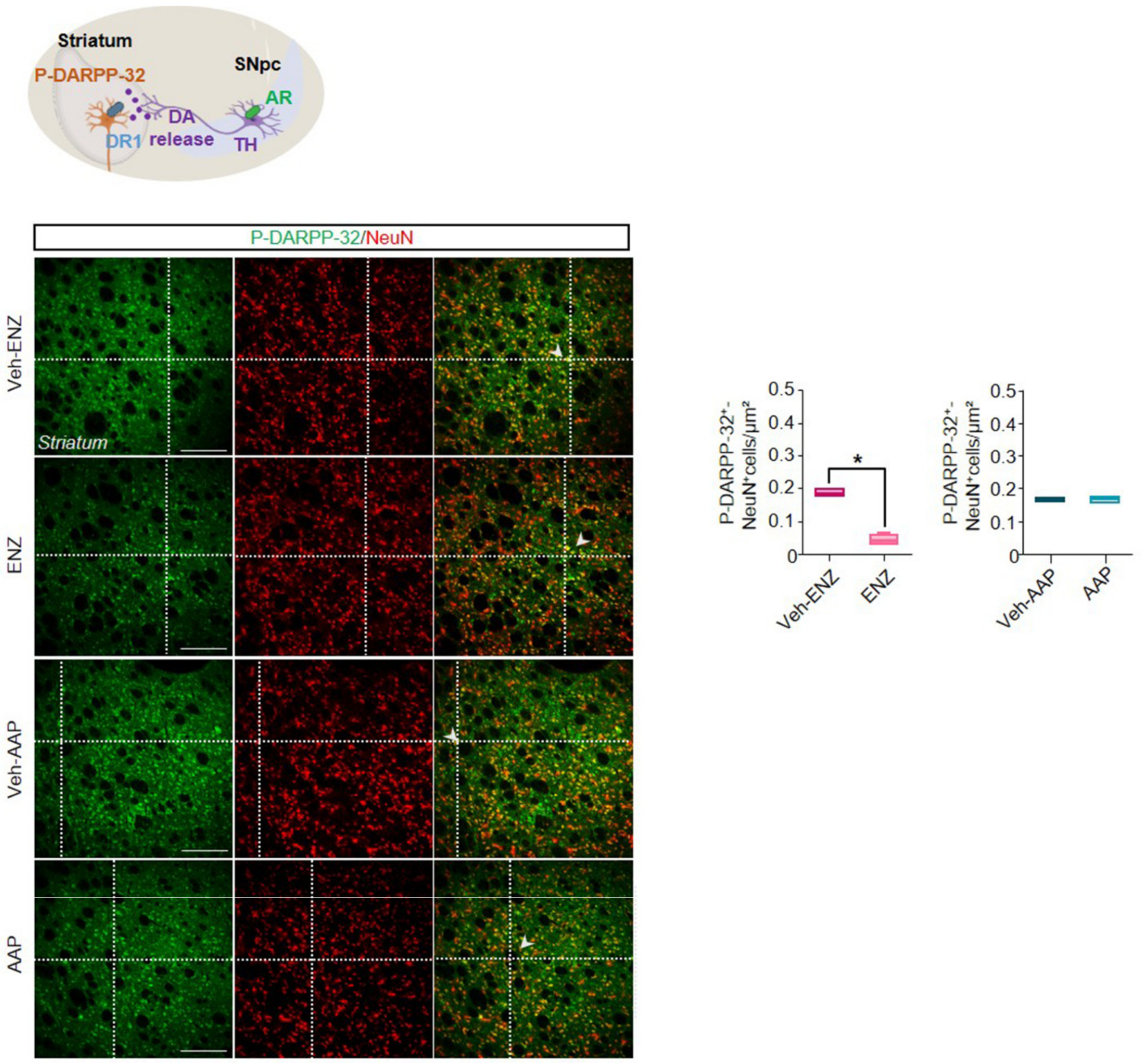


Nicola et al., S2 Fig

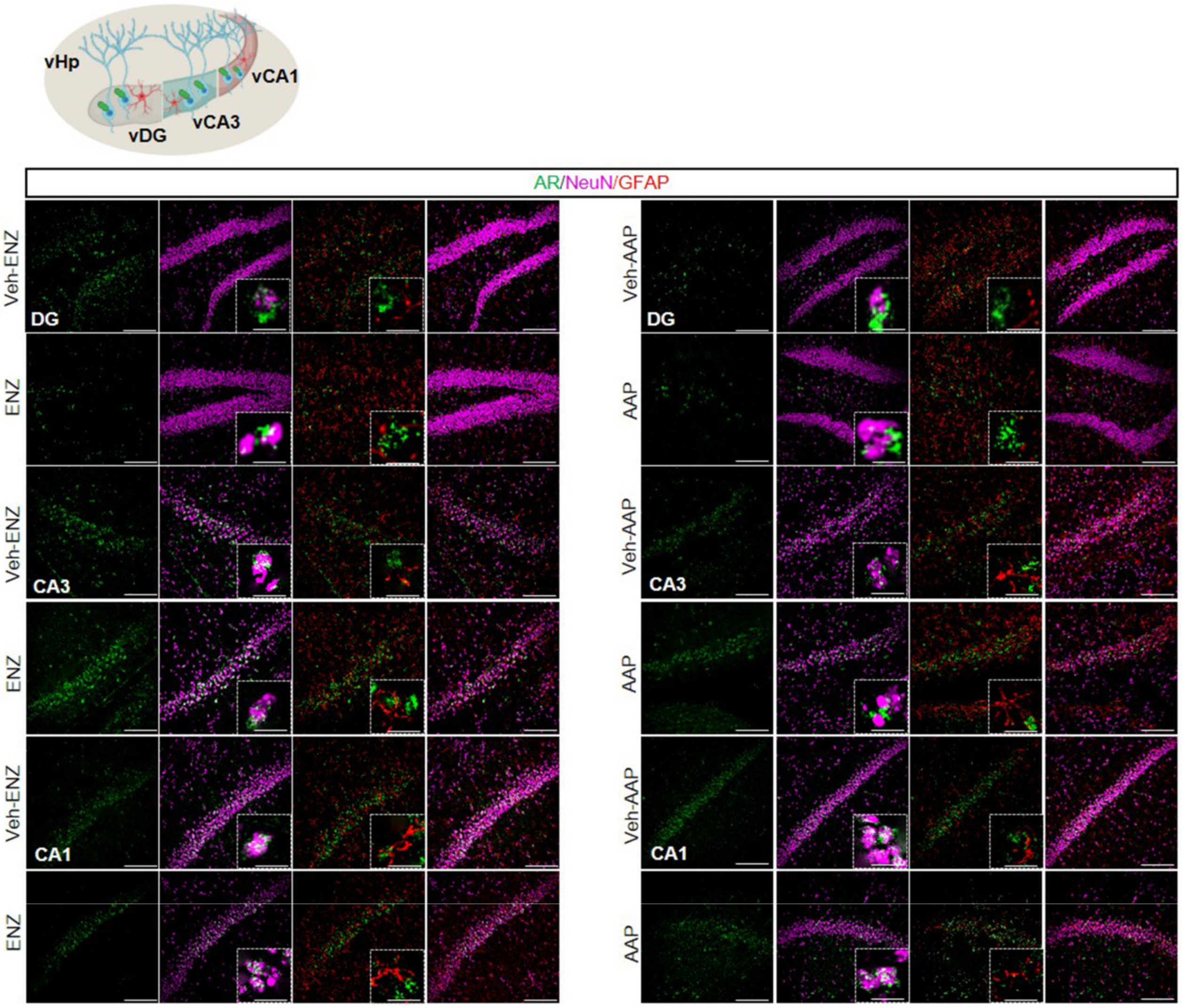


Nicola et al., S3 Fig

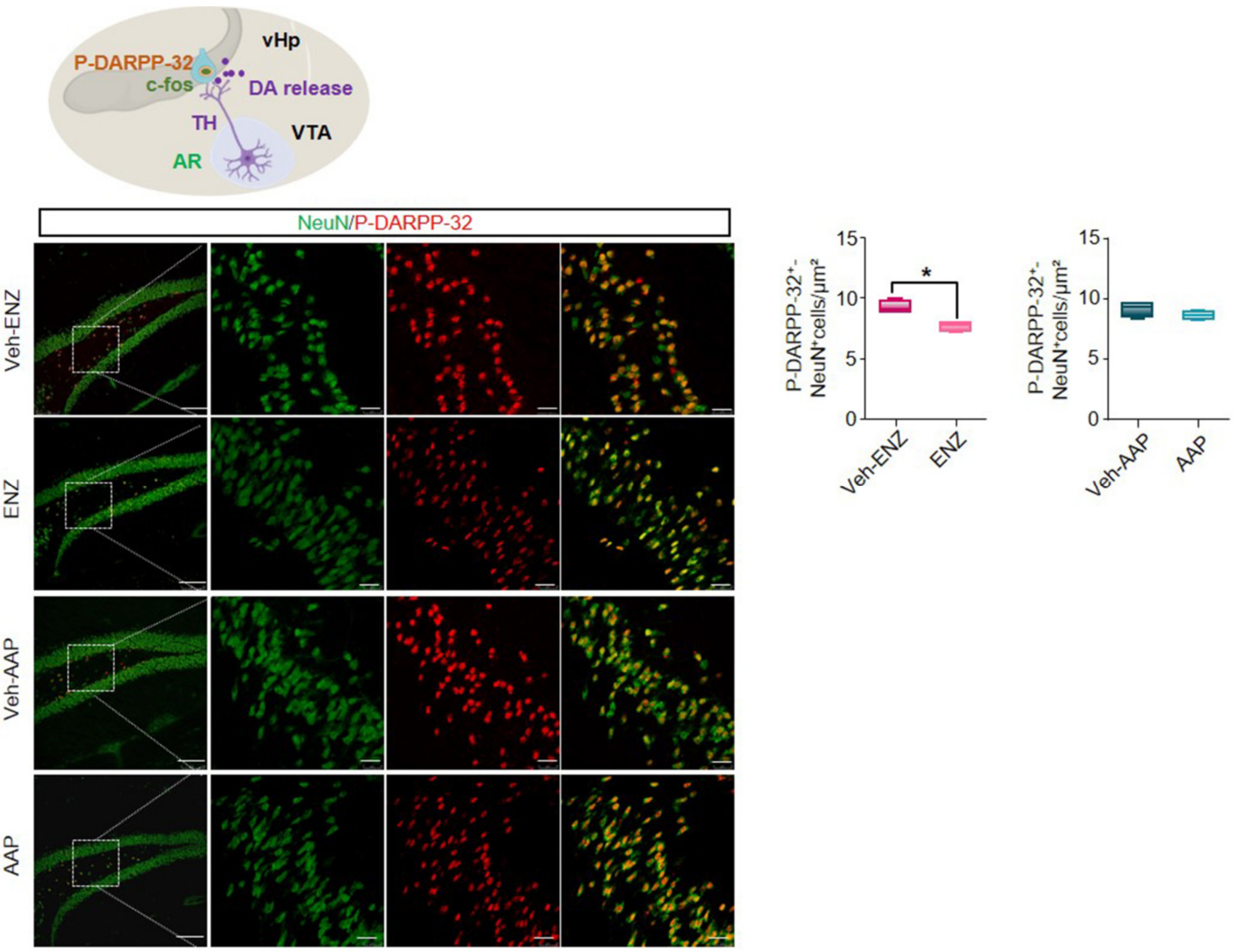


Nicola et al., S4 Fig

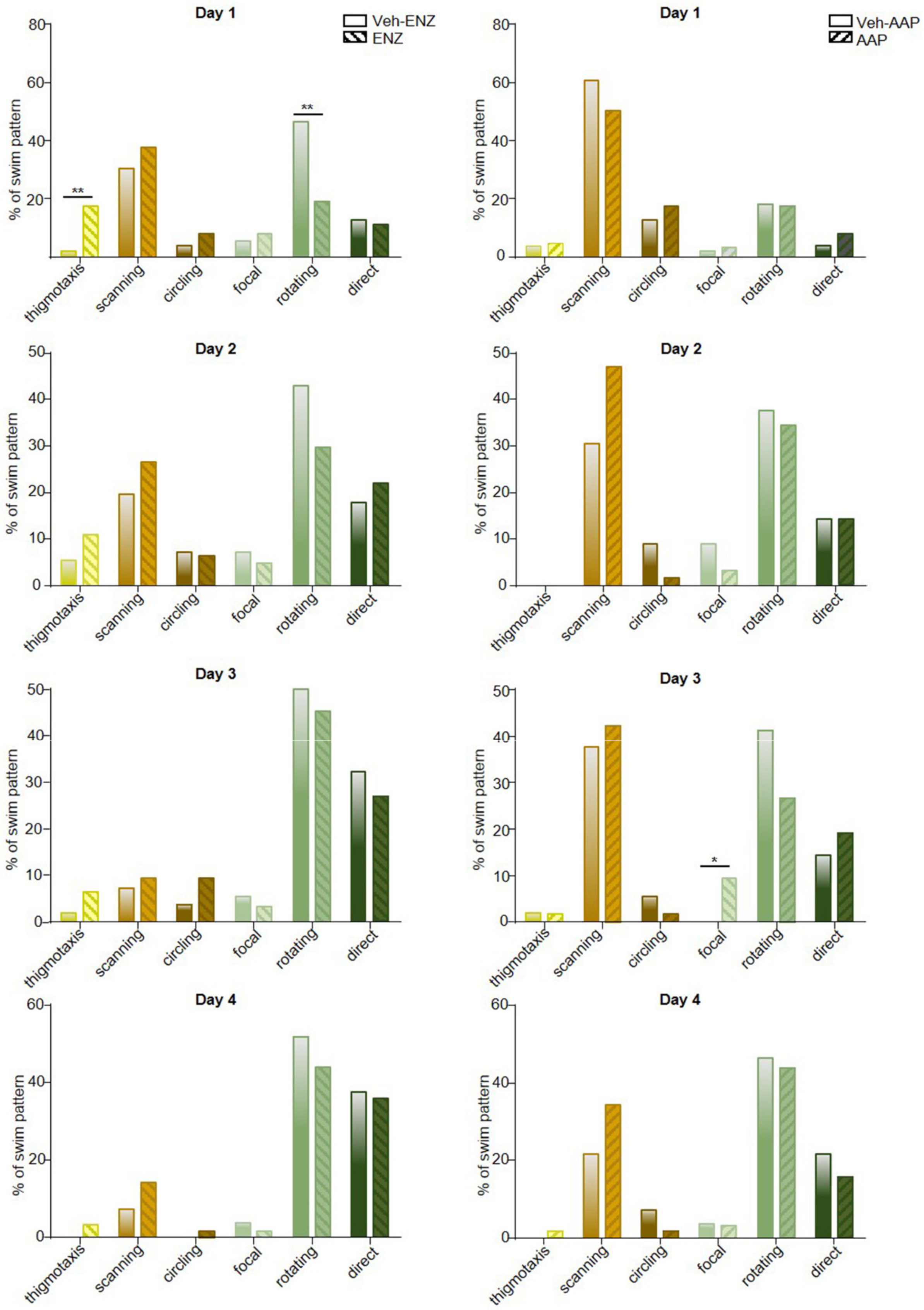


Nicola et al., S5 Fig

**A**

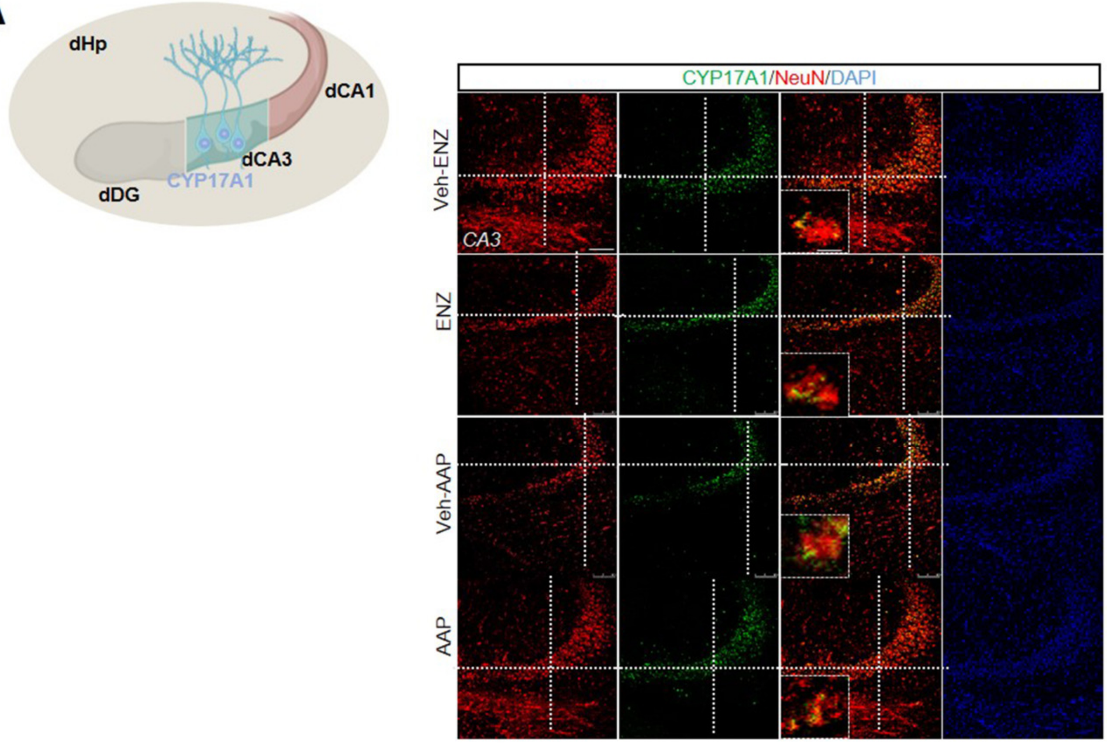
dHp
dCA1
dCA3
dDG
CYP17A1
CYP17A1/NeuN/DAPI
Veh-ENZ
CA3
ENZ
Veh-AAP
AAP

**B**

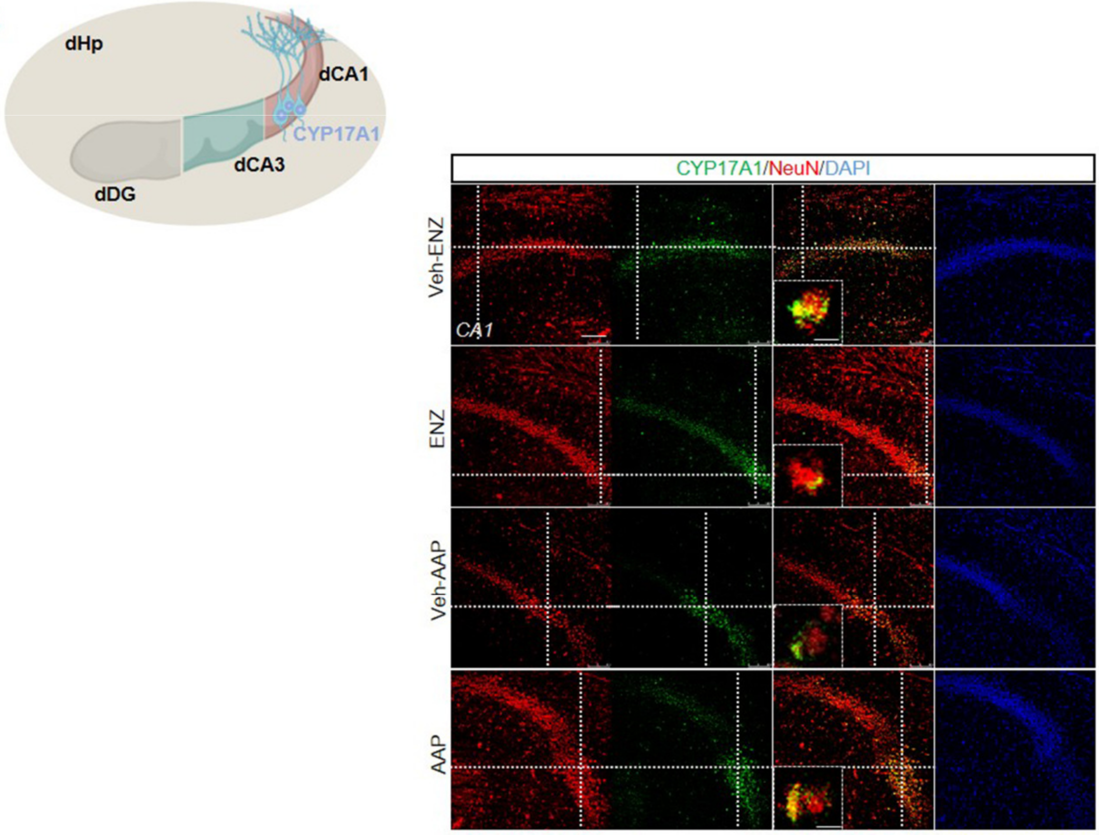
dHp
dCA1
CYP17A1
dCA3
dDG
CYP17A1/NeuN/DAPI
Veh-ENZ
CA1
ENZ
Veh-AAP
AAP

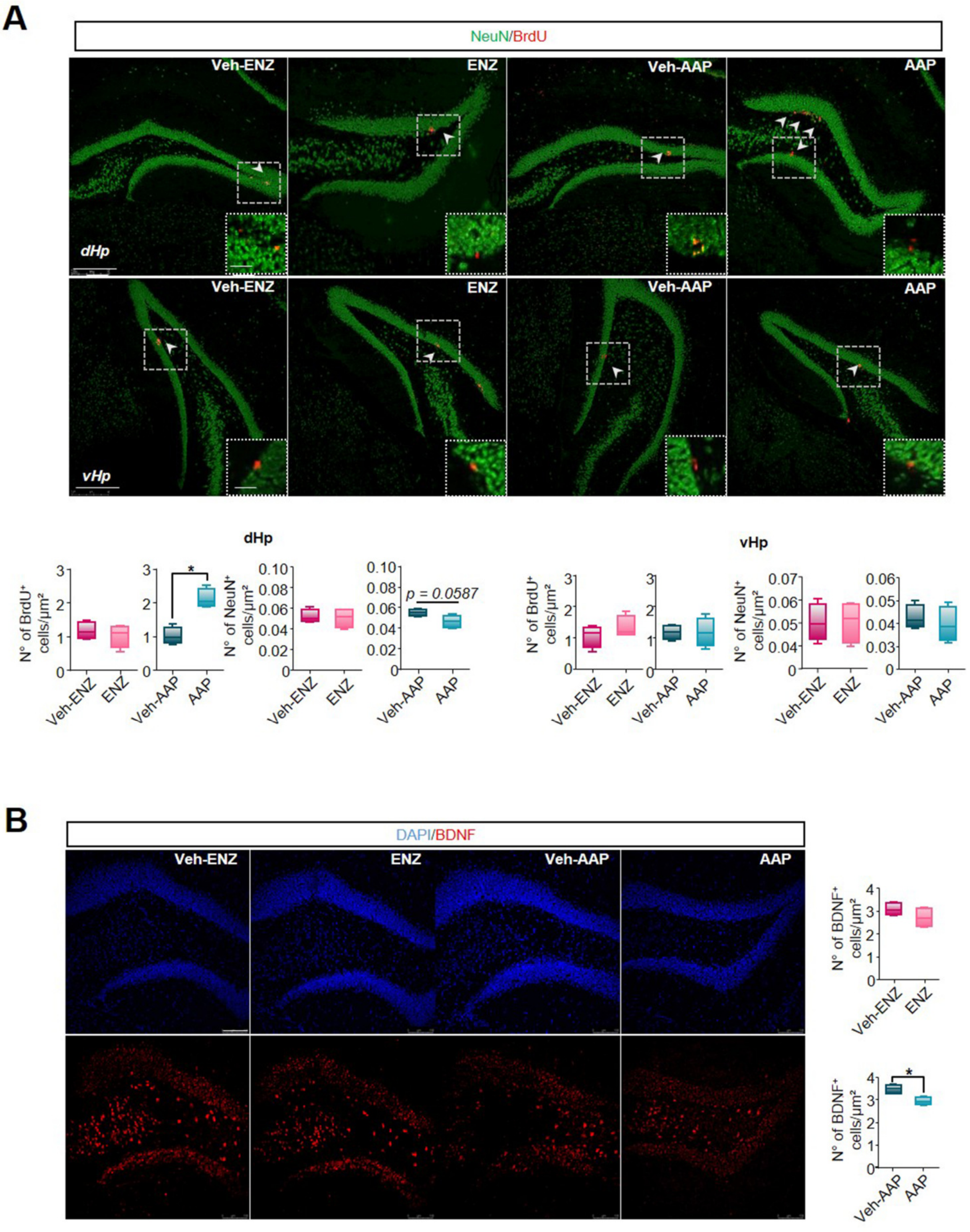


Nicola et al., S7 Fig
